# Transient frequency preference responses in cell signaling systems

**DOI:** 10.1101/2024.01.09.574874

**Authors:** Candela Lucia Szischik, Juliana Reves Szemere, Rocío Balderrama, Constanza Sánchez de la Vega, Alejandra C. Ventura

## Abstract

Ligand-receptor systems, covalent modification cycles, and transcriptional networks are the fundamental components of cell signaling and gene expression systems. While their behavior in reaching a steady state regime under step-like stimulation is well understood, their response under repetitive stimulation, particularly at early time stages is poorly characterized. This is despite the fact that early-stage responses to external inputs are arguably as informative as late-stage ones.

In simple systems, a periodic stimulation elicits an initial transient response, followed by periodic behavior. Transient responses are relevant when the stimulation has a limited time span, or when the stimulated component’s timescale is slow as compared to the timescales of the downstream processes, in which case these fast processes may be capturing only those transients. In this study, we analyze the frequency response of simple motifs at different time stages. We use dose-conserved pulsatile input signals, meaning that the amplitude or the duration of the pulses varies along with frequency to conserve input dose, and consider different metrics versus frequency curves.

We show that in ligand-receptor systems, there is a frequency preference response (band-pass filter) in some specific metrics during the transient stages, which is not present in the periodic regime. We suggest this is a general system-level mechanism that cells may use to filter input signals that have consequences for higher order circuits.

Additionally, we evaluate how the described behavior in isolated motifs is reflected in similar types of responses in cascades and pathways of which they are a part. Our studies suggest that transient frequency preferences are important dynamic features of cell signaling and gene expression systems, which have been overlooked.

## Introduction

### Cells responses and repetitive inputs

Cells can sense and respond to their environment through interconnected signaling pathways. These pathways transduce extracellular signals into gene expression patterns, ultimately shaping the cell’s behavior. The regulation of these cellular processes frequently relies on signals whose activity changes over time, often in a repetitive manner. Repetitive signals can originate from both external and internal sources (Chappell et al., 2003; De Pittà et al., 2009; Dunant et al., 1974; Skupin et al., 2010; Smedler & Uhlén, 2014; Thurley et al., 2014; Tomida et al., 2015), take the form of either periodic variations or irregular ones (e.g., quasi-periodic, stochastic), and play an important role in the precise control of cellular responses to environmental cues (Levine et al., 2013). The ability of repetitive signals to provide a more precise control over cellular responses can be particularly advantageous in noisy cellular environments. It is therefore crucial to determine which specific signal properties can optimize the activation of a particular target. This provides insights into the design principles of cellular signaling systems and the robustness of their responses to external perturbations such as environmental fluctuations.

### Examples of periodic inputs

Periodic signals can originate from both external and internal sources. The dynamic cellular environment often releases physiological ligands in pulsatile bursts with varying timescales, such as acetylcholine (Dunant et al., 1974), glutamate (De Pittà et al., 2009), and GnRH (Gonadotropin-releasing hormone) (Chappell et al., 2003), which can span from seconds to hours. In some cases, constant external stimulation can activate a signaling pathway that works as an oscillator, generating periodic inputs for downstream signaling modules. For example, oscillations in p38 MAP kinase activity are induced by continuous IL-1b stimulation and are necessary for the efficient expression of pro-inflammatory genes such as IL-6, IL-8, and COX-2 (Tomida et al., 2015). Calcium concentration is another example of a pulsatile signal, acting as a second messenger in diverse cellular processes and regulating targets based on its pulsatile dynamics (Skupin et al., 2010; Smedler & Uhlén, 2014; Thurley et al., 2014). Understanding the properties that determine the optimal activating signal for a specific target is essential to comprehend the regulatory repertoire of the cell in response to pulsatile signals.

### Different properties of a periodic or pulsatile input

Within repetitive inputs, we focus on periodic ones, and within this group we mainly consider pulsatile signals. These signals are defined by several key attributes, including pulse amplitude, duration, the time interval between consecutive pulses and frequency. While constant signals primarily regulate cellular responses through amplitude modulation, where the output is determined solely by the strength of the input, pulsatile signals add additional layers of modulation as the attributes mentioned above may also impact the cellular responses (Martinez-Corral & Garcia-Ojalvo, 2017). A significant type of modulation is the frequency dependence of the cellular responses.

### Classification of frequency responses and experimental examples

Biological systems can exhibit a range of frequency-dependent responses to periodic signals, which can be classified into all-, low-, band- and high-pass filters. All-pass filters describe situations where the response is not affected by the frequency of the stimulus. One example is the NFAT-mediated transcriptional regulation, which shows no significant difference in reporter expression across calcium frequencies (Hannanta-Anan & Chow, 2016). Low-pass filters integrate the signal when it changes rapidly and follow it more faithfully when it changes slowly. The high-osmolarity glycerol (HOG) signaling pathway in *Saccharomyces cerevisiae* and the nitrogen assimilation control network in *Escherichia coli* are examples of such systems (Hersen et al., 2008; Jiang et al., 2011). Band-pass filters show an optimal (or preferred) response (resonance) for an intermediate range of input frequencies. For instance, osmo-adaptation in Saccharomyces cerevisiae exhibits frequency selectivity (Mettetal et al., 2008). For another example, (Mitchell et al., 2015) found that an intermediate frequency causes slow proliferation in yeast subjected to oscillatory osmotic stress. Such resonant responses can be essential for biological systems to selectively respond to specific signals and stimuli. Finally, high-pass filters behave opposite to low-pass filters in the frequency domain.

### The response of interest

The frequency-dependent behavior of the response signals to periodic inputs is attribute-dependent. To assess whether a signaling or gene expression system exhibits band-pass filtering behavior, it must first be determined which attribute of the response is the appropriate one to be evaluated. It is conceivable that some response attributes may exhibit resonance while others do not. For example, Hersen et al. (Hersen et al., 2008) measured the amplitude of the oscillating level of the third level of the MAPK cascade to assess the frequency-dependent response. Hannanta-Anan and Chow (Hannanta-Anan & Chow, 2016) computed the accumulated integral in NFAT transcriptional regulation. Johnson and Toettcher (Johnson & Toettcher, 2019) also used this operation to study cell fate in *Drosophila* embryo. Another possible response is the output’s integral computed over one period of the input, as used in (Ryu et al., 2015) for the ERK activity in response to different pulse/pause regimes. Zi and collegues (Zi et al., 2010) quantified the signal transduction efficiency of Hog1 response to different scenarios of osmotic stress in *Saccharomyces cerevisiae* using the gain, defined as the integrated output divided by the integrated input. For yet another example, Pena & Rotstein (Pena & Rotstein, 2022) showed that two-dimensional linear models receiving pulsatile stimulation exhibit resonance in the impedance amplitude curve, but a low-pass filter in the peak-responses versus frequency curve.

### The time window where the response is calculated

Although often overlooked, the frequency-dependent response of a system to periodic inputs is time window-dependent. Specifically, this type of responses can be calculated over various temporal windows and there is no reason to believe knowledge of the response over one (e.g., steady-state or quasi-steady state) will be predictive of the other (e.g., transient). For example, one can consider an early temporal window (transient regime), a late temporal window (periodic behavior if it exists) or a global window that encompasses both the transient and periodic regimes. Marhl et al. (Marhl et al., 2006) provided evidence that simple signaling motifs, which would be expected to act as low-pass filters, behave in fact as band-pass filters if the input oscillations are time-limited. Signals such as Ca^2+^ oscillations are time-limited and activate processes that are also time-limited, thus preventing the system from achieving stationary periodic behavior.

Even when periodic stimulation is prolonged, signaling components may still be functioning at pre-equilibrium. For instance, in *Saccharomyces cerevisiae*’s mating response, the signaling component that processes a stimulus operates on a slower timescale than its downstream processes. When the concentration is close to the dissociation constant, binding takes around 20 minutes to achieve 90% of the equilibrium level. These dynamics are not only slower than the cell division cycle (90 minutes) but they are also slower than the pheromone-dependent activation of the mitogen-activated protein kinase (MAPK) Fus3, which takes about 2 to 5 minutes to attain steady-state levels (Ventura et al., 2014). As a result, it is reasonable to conclude that the machinery downstream of the pheromone receptor must rely on transient binding information to function effectively. An implication of the two situations described above (signals are limited in time or the timescales of the downstream signaling component are fast as compared to the receiving component) is that pathways use information coming from the transients. In most of the articles where the frequency-dependent response is studied, the initial transient behavior (first portion of the response) is discarded.

### Summary of what was done in the article

In this article we study the ability of simple cell signaling motifs to produce transient frequency preference responses (band-pass filters or resonance) to dose-conserved pulsatile stimulation. We focus on arguably the simplest motif where the system exhibits sustained (monotonic) activation in response to constant inputs and are steady-state low-pass filters (in response to oscillatory inputs, particularly pulsatile inputs). We leave out of this study systems that produce sustained oscillations (limit cycles) and transient activation (adaptation, overshoot) in response to constant inputs. These two groups are expected to show preferred frequency responses in the stationary regime. Biologically, within the motifs of interest, we first consider a ligand-receptor reaction and a synthesis-degradation signaling component (Heinrich et al., 2002; Krivan et al., 2002; Tchourine et al., 2014). We then extend this system to a cascade by coupling the simple signaling components with feedforward connections. The dose-conservation scheme we use (Fig. 1B) consists of adjusting the pulse amplitude or duration to compensate for change in the pulse-frequency and therefore avoid a dose increase with increasing frequencies (Fletcher et al., 2014). We measure the systems’ response to pulsatile inputs by computing the average response over a sliding window of width equal to the input period and over an expanding window starting at zero (Fig. 2A). These two observables provide complementary information about the transient vs. steady-state nature of the preferred frequency response to pulsatile inputs and the transition of these responses as time progresses. We divide the systems’ response to pulsatile inputs into two regimes (Fig. 1A): transient and periodic. While strictly speaking periodic behavior occurs only in the stationary regime (i.e., it is reached as time approaches infinity), for simplicity in the description, in this article we refer to the asymptotically periodic behavior approximating the stationary periodic behavior as periodic.

**Figure 1.**
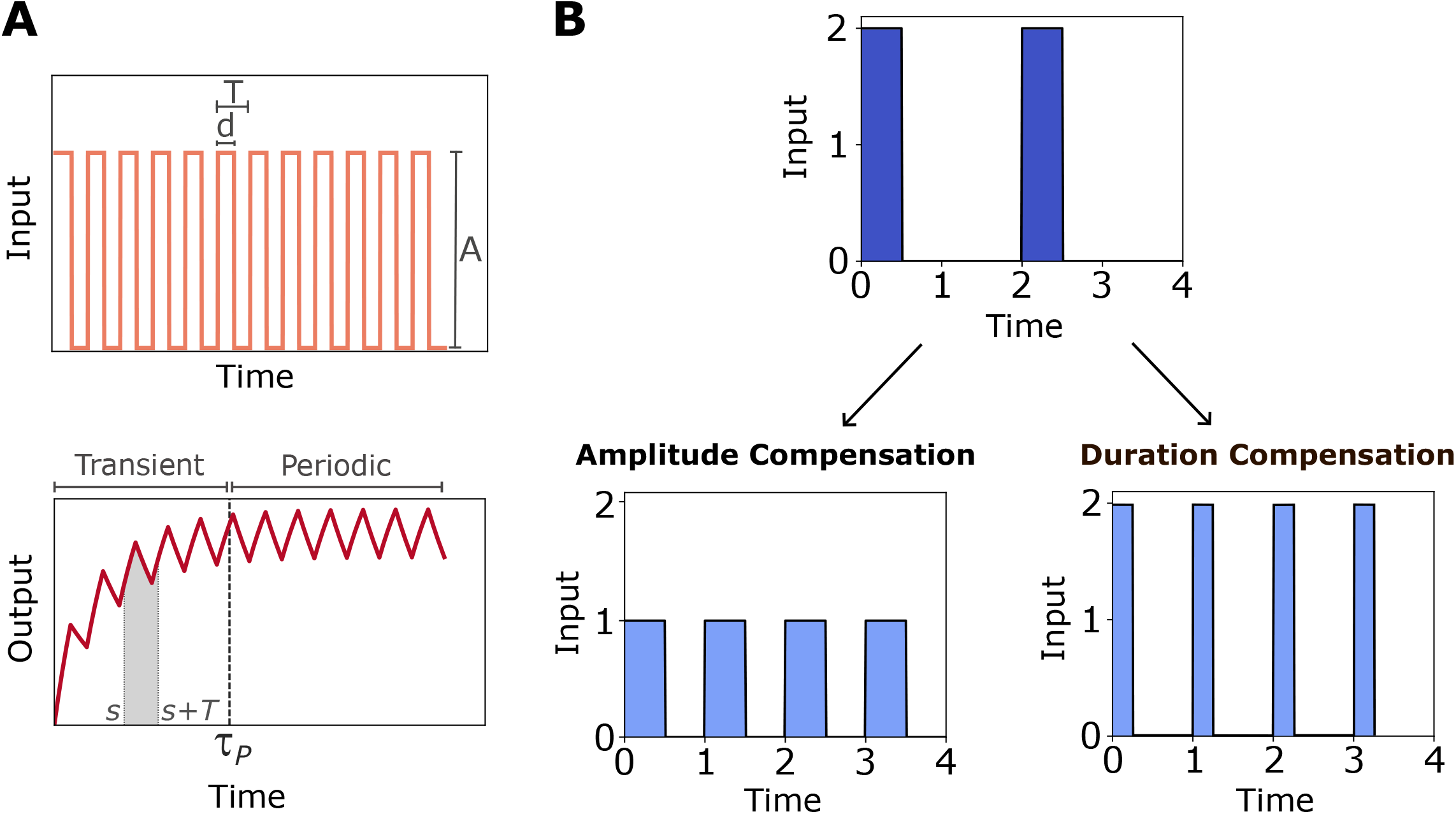
Ligand-receptor system’s response to periodic stimulation. **A**. Upper panel: system’s input as a train of square pulses, where *A* is the pulse amplitude, *d* is the duration of each pulse and *T* is the period. The lower panel contains an example of an output to this periodic input, the scaled concentration of complex *y*(*t*), as a function of time. The time it takes to the system to exhibit periodic oscillations is defined as the characteristic time τ_*p*_, thus defining two dynamical stages: transient and periodic regimes. An example of a temporal window, between *s* and *s+T* is shown in grey. **B**. Dose conservation protocols: by amplitude and duration compensation (left and right panel, respectively).

**Figure 2.**
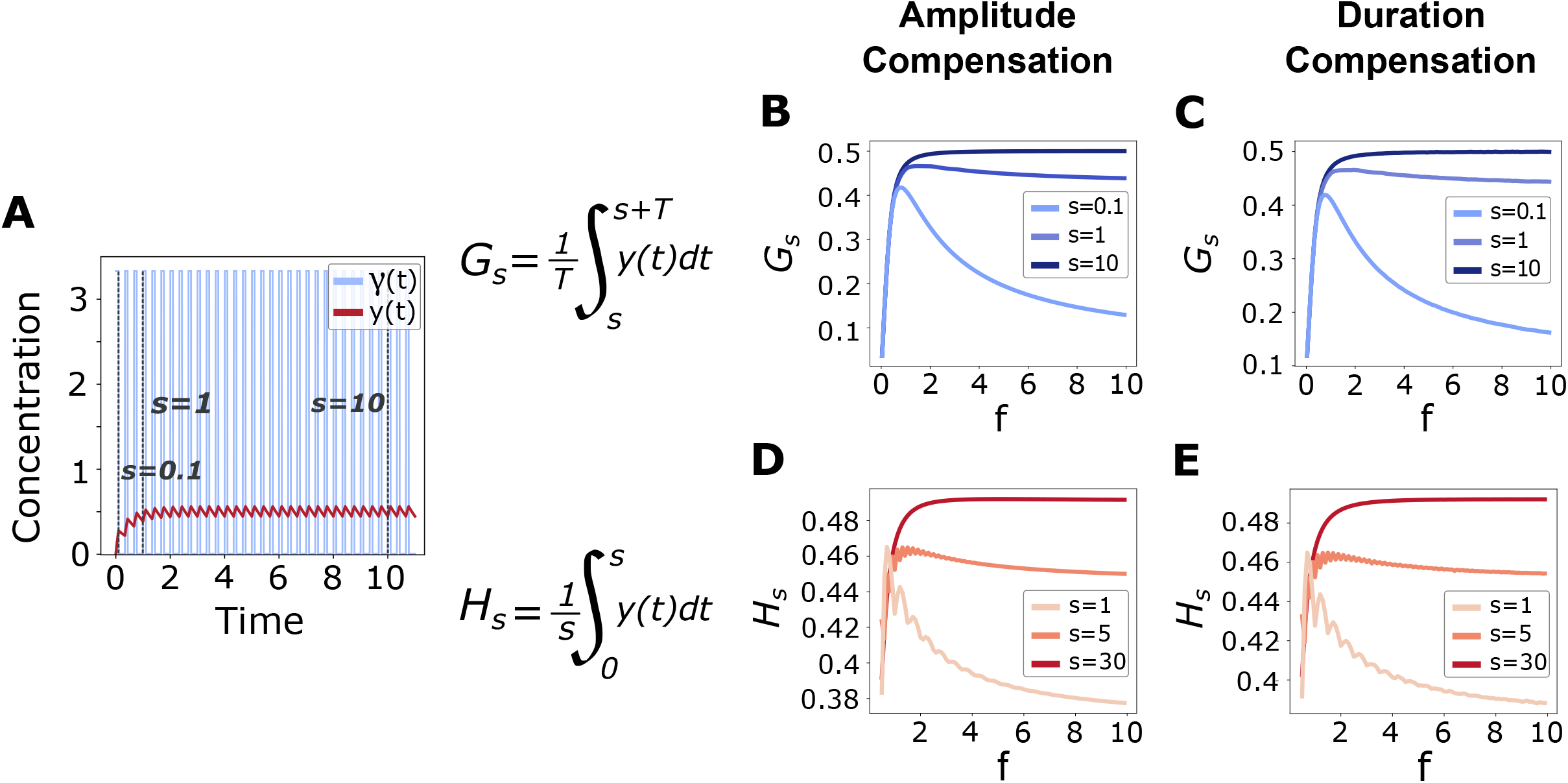
Transient frequency preference response for the ligand-receptor system. **A**. Input concentration -free ligand- (light blue) with parameters *T* = 1/_3_, *A = 3*._3_, *d* = 0.1 and the system’
ss response to this input -ligand-receptor complex-in red as a function of time. Dotted vertical lines indicate different values of *s* selected to compute the observables in panels **B-C** (*s =0*.*1, s=1* and *s* = 10). **B** and **C**. *G*_*s*_ versus frequency for the two dose conservation protocols: by amplitude compensation in **B** (*T*_0_ *= 1, A*_0_ = 10, *d* = 0.1, thus < *γ >* = 1) and by duration in **C** (*T*_0_ *= 1, A* = 10, *d*_0_ = 0.1, thus < *γ > =1*). **D** and **E**. *H*_*s*_ versus frequency for the two dose conservation protocols: in **D** by amplitude compensation and in **E** by varying the duration of the pulses.

The major contribution of this work lies in the study of frequency responses considering a temporal window during which the output of the stimulated system is in a transient phase. We found that in several simple systems there is a frequency preference response in some specific metrics during the transient stages, which is not present in the periodic ones. Additionally, we evaluated how the described behavior in isolated motifs is reflected in similar types of responses in cascades and pathways of which they are a part. Our studies suggest that transient frequency preferences are important dynamic features of cell signaling and gene expression systems, which have been overlooked.

## Results

### 1. Ligand-receptor reaction: introducing the notion of transient frequency preference response

One of the simplest binding models is a receptor ***R*** that binds a ligand ***L*** forming a complex ***C***. It is described by the following reversible reaction:

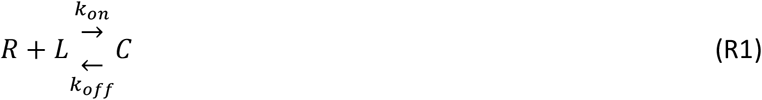

where *k*_*on*_ is the association rate constant and *k*_*off*_ is the dissociation rate constant. Assuming mass action kinetics, the total amount of receptor is constant (***R***_***T***_ *=* ***R*** *+* ***C***) and the concentration of free ligand is not depleted significantly by the binding reaction (***L*** *∼* ***L***_***T***_), the dynamics of the complex ***C*** can be described by the following equation:

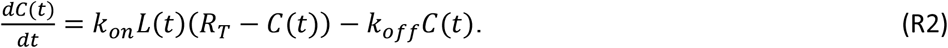

We define the following variables to nondimensionalize eq. R2:

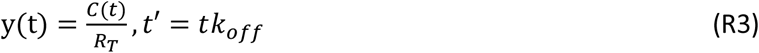

Substituting into (R3) yields to:

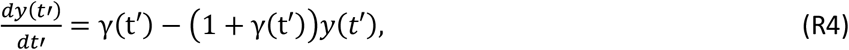

where *γ*(*t’*) *= L*(*t’*/*k*_off_)/*K* is the amount of ligand relative to the dissociation constant for the binding-unbinding reaction *K = k*_*off*_ /*k*_*on*_. From now on we refer to the dimensionless time *t’* as *t* to simplify notation.

Note that Eq. R4 has no parameters except for the ones that correspond to the stimulus, meaning that this model could describe the dynamics of any ligand-receptor reaction. Because of the nondimensionalization, the variable y is bounded, as *C ≤ R*_*T*_, then 0 *≤ y* ≤ 1 (see Supplementary Information, section 1.1.2). When *γ* is constant, the time the system takes to reach to the steady state is: τ = 1/(1 *+ γ*) (see Supplementary Information, section 1.1.2 and (Ventura et al., 2014)). This is the characteristic time for the ligand-binding reaction and is relevant for future analysis in this section.

The ligand-receptor system is stimulated with a train of square pulses of ligand, as shown in Fig. 1. Specifically, we consider *γ* to be the *T*-periodic function defined in [0, *T*] as follows:

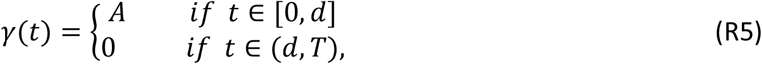

We define < *γ >*, the mean dose of ligand in a period (*T*), as

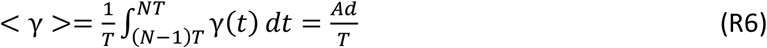

Following (Fletcher et al., 2014), when the frequency of the input is modified, the total dose of the stimulus is conserved using two different protocols based on modifying either the amplitude or the duration of the pulse (Fig. 1). For the amplitude compensation protocol (fixed duration), starting from reference stimulus attributes of *A*_0_ (amplitude), *d*_0_ (duration) and *T*_0_ (period), if the period is modified to *T*, then the amplitude is adjusted according to 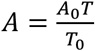. For the duration compensation protocol, the pulse duration is adjusted to *d = d*_0_*T*/*T*_0_. In both cases, the mean dose per period < *γ >* is *T* -independent.

It can be shown that the system described by Eq. R4 always oscillates in response to a periodic stimulus, and that if the periodic stimulus is a train of square pulses, it takes a time *τ*_*p*_ to reach an almost periodic solution (see Supplementary Information, section 1.1.1). Therefore, we can distinguish two regimes in the output: a transient regime (*t < τ*_*p*_) and a periodic regime (*t > τ*_*p*_). In the latter one we find almost periodic oscillations (technically, truly periodic oscillations are displayed only in the stationary regime as *t* approaches infinity). Moreover, in certain conditions τ_*p*_ can be bounded by the time *τ* it takes the system to reach the steady state when *γ* is constant (see Supplementary Information, section 1.1.2).

To study the frequency response of the ligand-receptor system, we computed two observables for each of the dose conservation protocols described above. The sliding window:

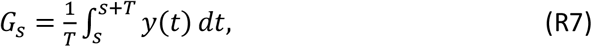

and the expanding window:

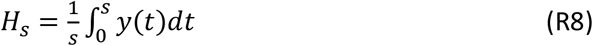

*G*_*s*_ is the output average over a period *T c*omputed between an initial time *s* to *s + T*, whereas *H*_*s*_ is the output average computed from 0 to a final time *s*. Both observables are bounded between 0 and 1, as the output *y* is also bounded between these values. As the parameter s is increased, the observables are computed over the different dynamical stages of the output and capture the response properties in the transient and steady-state regimes both locally and globally as well as the transitions between them.

Fig. 2A shows the results for a given reference input. The input parameters (*T*_0_ = 1, *A*_0_ = 10, *d*_0_ *= 0*.1) are such that the mean dose over a period is < *γ >*= 1. If we consider the equivalent constant input with the same mean dose, the time it takes to reach a steady state is 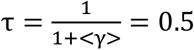. This characteristic time represents a good indicator of the transition between the transient to the periodic regimes in response to periodic inputs.

The procedure was repeated for the two dose compensation schemes and different values of *s*, going from a transient stage (*s* = 0.1) to a periodic one (*s* = 10). The frequency response of the ligand-receptor system is highly dependent on parameter *s* (the dynamical stage the system is facing). Local fluctuations are observed over the asymmetric bell-shaped curve of the observable *H*_*s*_ as a function of frequency. These oscillations are due to small variations in the mean dose over the time interval [0, *s*] where the observable is computed, since it may not contain a complete number of periods. If one period is contained only partially, its contribution to the integral could introduce variations in the observable. No local fluctuations are found in the observable *G*_*s*_, as the time window -from *s* to *s+T*-covers entire periods, therefore the mean dose per period is constant for all the frequencies and values of *s*.

Results in Fig. 2B to E show that if the observables are computed in a transient phase of the output (smaller values of *s*), then they exhibit a band-pass filter behavior. In contrast, for longer times (higher values of *s*) the observables exhibit a high-pass filter behavior (there is a frequency cut-off, from which the response is insensitive to the frequency). We refer to the former situations as a **transient frequency preference**. This behavior has proven to be robust since it has been observed in the four combinations of observables and input dose conservation protocols. Transient frequency preference also emerges from a sinusoidal stimulation (see Supplementary Information, section 4.1) and without dose conservation (see section 6). These results suggest that a signaling component, such as the ligand-receptor system, can send different messages, depending on the typical timescales of the downstream components receiving those messages.

In Fig. 3 we evaluate how the stimulation parameters -pulse duration *d*_0_, pulse amplitude *A*_0_ and mean input dose 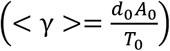-shape the frequency preference response. Once the reference parameters values are fixed, dose conservation by an amplitude compensation protocol was applied for all the frequencies considered. This study was performed for the sliding window *G*_*s*_ and for an initial time *s* such as the output is in the transient regime (*s* = 0.1). In order to quantify the parameters’ influence in the transient frequency preference response, we compute the preferred frequency *f*_*G,max*_, and its corresponding value in the curve *G*_*s*_, *G*_*max*_, versus dose. A similar study in the periodic regime is included in the Supplementary Information, section 4.2.1.

**Figure 3.**
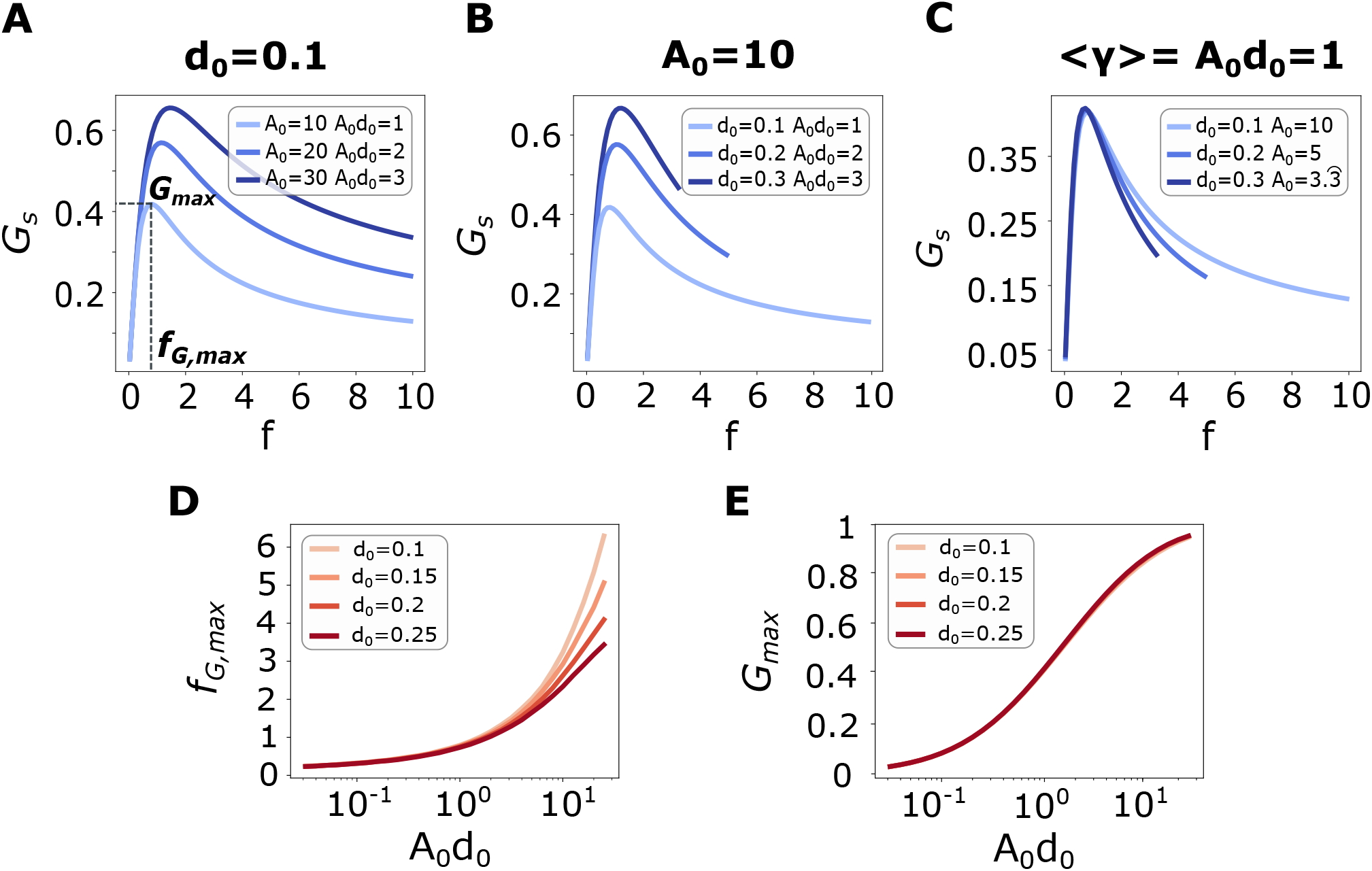
Input parameters’ influence on the transient frequency preference response for the ligand-receptor system: **A-C**. Sliding window *G*_*s*_ as a function of frequency, measured at a transient time *s* = 0.1. Dose was conserved by varying the pulses’ amplitude, for all simulations *T*_0_ = 1. **A**. Fixed reference pulse duration *d*_0_ *= 0*.1 and different reference amplitudes *A*_0_, so that the mean dose < *γ >*= *A*_0_*d*_0_ increases from 1 to 3. **B**. Fixed amplitude *A*_0_ = 10 and different reference pulse durations, so that the mean dose < *γ >*= *A*_0_*d*_0_ *i*ncreases from 1 to 3. **C**. Fixed dose for the three curves, varying *d*_0_ and *A*_0_. Each curve was simulated until the maximum frequency possible, given by *f*_*cut*_ = 1/*d*_0_. **Transient frequency preference dependence on the mean dose: D-E. D**. Preferred frequency for different frequency curves of the observable *G*_*s*_. *f*_*G,max*_ as a function of the mean dose *A*_0_*d*_0_. ***E***. Maximum value of the observable *G*_*s*_, *G*_*max*_, as a function of the mean dose *A*_0_*d*_0_. The duration *d*_0_ *w*as fixed as the amplitude *A*_0_ varies. Different durations *d*_0_ are indicated in colors.

In Fig. 3A, we consider *T*_0_ = 1, a fixed duration *d*_0_ = 0.1 and varying values of *A*_0_, resulting in different mean input doses < *γ > = A*_0_*d*_0_. As *A*_0_ increases, the preferred frequency *f*_*G,max*_ shifts towards higher values and *G*_*max*_ also increases. A similar result is found by fixing the reference amplitude *A*_0_ and increasing the pulse duration *d*_0_ (Fig. 3B). In Fig. 3C, both *d*_0_ and *A*_0_ are varied while keeping the mean input *d*_0_*A*_0_ constant. Only in this case, *f*_*G,max*_ and *G*_*max*_ remain constant, suggesting that the preferred frequency and maximum value for the sliding window are controlled by the mean input dose. In Fig. 3D and E we explore the influence of the mean input dose on the frequency preference response. The preferred frequency *f*_*G,max*_ and the maximum value *G*_*max*_ are plotted as functions of the mean dose. Similar results are found using the duration compensation scheme for dose conservation, shown in Supplementary Information, section 4.2.2.

The preferred frequency *f*_*G,max*_ *v*aries with the mean input dose in a similar way for the different values of *d*_0_. Although there are some differences in the curves for higher doses, they are not significant compared to the variations in dose (Fig. 3D). Note that the scale of the x-axis (mean dose) is logarithmic, while the y-axis scale (*f*_*G,max*_) is linear. Regarding the maximum value *G*_*max*_ (Fig. 3E), its dependence with the dose is not modified by varying *d*_0_.

Together, these studies support the underlying hypothesis that the fundamental parameter to understand the transient frequency response in a ligand-receptor system is the input mean dose.

### 2. Transient frequency preference response in signaling components with a simplified linear description

In this section we follow Fletcher et al. (Fletcher et al., 2014) and extend our study to a signaling component with a slightly different description from the one in the previous section. The model is described by the following equation:

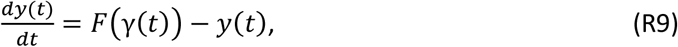

where the variable *y*(t) is the dimensionless concentration of a given molecule, the time is scaled by a reference time and γ(*t*) is the input.

Equation R9 is a simplified model that could represent several biological systems, such as synthesis and degradation of proteins. The model is linear in the variable *y* and includes a function *F t*hat processes the input γ(*t*). The function *F* can be thought of as a rapid equilibrium approximation to fast signaling dynamics that occur prior to the production of *y* (Fletcher et al., 2014).

For a constant input *γ*(*t*) *= A, y* reaches the steady state *F*(*A*), so *F* represents the input-output curve for constant stimulation. For a square-pulse input, *y* rises towards the steady state *F*(*A*) during the pulse and decays towards zero after the pulse ends. For an input consisting of a train of square pulses, there is a steady-state periodic response resulting from a sequential combination of the increase and decrease phases described above (see Supplementary Information, section 1.2).

In this section we focus on the effect of the nonlinearity of *F* on the frequency response curves *G*_*s*_ of the simplified linear model. Our goal is to identify the choices of the nonlinear production rate *F* that result in a band-pass filter for the sliding window *G*_*s*_ and whether band-pass filters arise during the transient regime, the periodic regime or both. Based on previous research (Fletcher et al., 2014), our study considers three types of monotonic increasing functions *F*: convex, concave, and sigmoidal functions. A linear function lies in the limit between convex and concave functions. The study conducted by Fletcher and co-authors focused solely on the periodic response. We extend this study to incorporate the transient response. We examine the observable *G*_*s*_ as a function of the input frequency, considering an amplitude compensation scheme to maintain dose conservation.

Fig. 4 shows the graphs of *G*_*s*_ (Fig. 4B) for functions *F* with representative shapes (Fig. 4A) and for different values of *s*. These results are supported by analytical calculations in the Supplementary Information, section 2. From our numerical and analytical studies, we conclude that if *F* is a linear or convex function (Fig. 4, rows 1 and 2), *G*_*s*_ is a low-pass filter both for the transient and periodic regimes. If *F* is a concave function, then *G*_*s*_ is a high-pass filter for the periodic regime. For the transient regime, *G*_*s*_ can be a low- or a band-pass filter depending on the choice of the function *F*. We present two examples in Fig. 4 (row 3). The left panel corresponds to the function 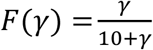, having a transient frequency preference, and the right panel corresponds to the function 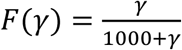, having a low-pass filter behaviour. Finally, if *F* is a sigmoidal function, the behaviour of *G*_*s*_ also depends on the choice of *F*. We present two examples in Fig. 4 (row 4). For *F*(*γ*) *= γ*^2^/(1 *+ γ*^2^) (left), the response has a frequency preference both in the transient and in the periodic regime, for *F*(*γ*) *= γ*^2^/(1000 *+ γ*^2^) (right), the response behaves as a low-pass filter.

**Figure 4.**
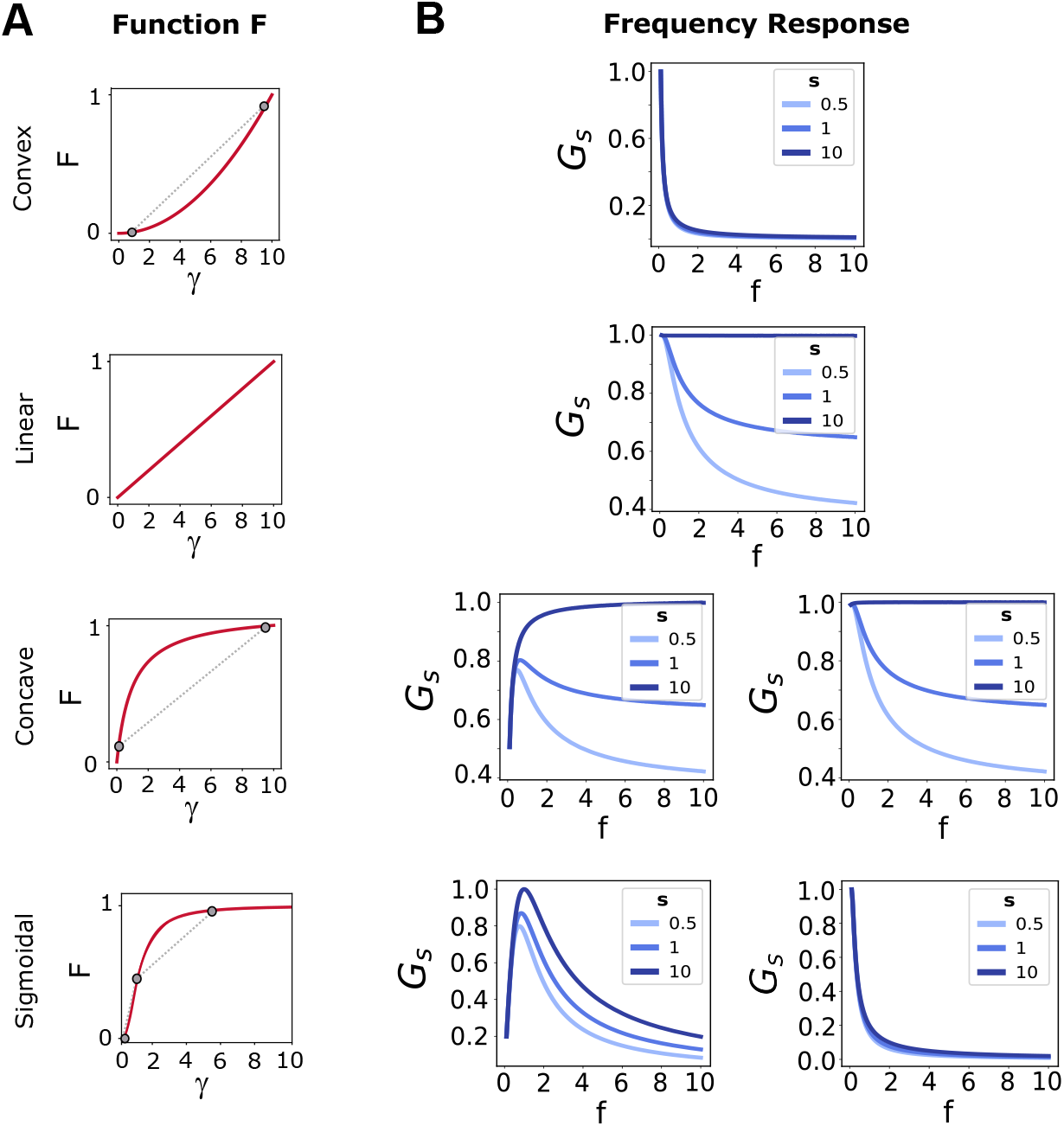
The frequency response for a signaling component with a simplified linear description depends on the concavity of the function *F*(γ). The system is stimulated with a periodic train of square pulses, using an amplitude compensation protocol to conserve input dose. Input parameters: *A*_0_ *= 1, d*_0_ *= 1, T*_0_ = 1. The observable computed is the sliding window *G*_*s*_, as described in Methods, normalized by its maximum value. **A**. Examples for the four possible concavities for function *F*(*γ*). **B**. Frequency response for the four concavities (rows) for different values of *s*, earlier times in lighter lines. For concave and sigmoidal functions, different responses are achieved for different parameter values. The functions selected for this figure are: convex *F = γ*^2^, linear *F = γ*, concaves *F = F = γ*/(10 *+ γ*) (left) and *F = F = γ*/(1000 *+ γ*) (right) and sigmoidals *F = γ*^2^/(1 *+ γ*^2^) (left) and *F = F = γ*^2^/(100 *+ γ*^2^) (right).

Results for the observable *H*_*s*_ are shown in Supplementary Information, section 5. For this observable, there is a minimum frequency given by 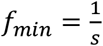 to have at least one oscillation in the time window. Thus, for shorter initial times the range of frequencies is narrower.

In summary, consistently with the results of Fletcher et al. (Fletcher et al., 2014) our numerical and analytical results show that only models with sigmoidal functions *F* can produce band-pass filters in the periodic regime. However, models with both sigmoidal and concave functions *F* can produce band-pass filters in the transient regime. These results are not true for all sigmoidal and concave functions *F*, but the existence of transient band-pass filters depend on the choice of the parameters of *F*.

### 3. Coupled signaling components without feedback connections

In the preceding sections, we focused on examining the frequency preference responses in elementary signaling components, such as ligand-receptor reactions and linear signaling units with additive nonlinear processing of pulsatile inputs. Our findings revealed that both systems exhibited frequency selectivity under some conditions when the observables *G*_*s*_ and *H*_*s*_ were evaluated in the transient regime (see also examples 2.1, 2.2 and 2.3 in Supplementary Information). To further explore this phenomenon, we now turn our attention to a more intricate scenario: a cascade consisting of two signaling components interconnected by a feedforward connection. The key issue we aim to address in this section is whether the transient frequency preference observed in the earlier systems is transmitted through the cascade. By investigating this more complex configuration, we seek to gain insights into the behavior and potential amplification or attenuation of the transient frequency preference within the cascade structure.

The cascade system is described by:

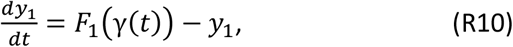

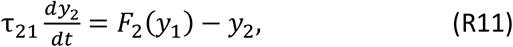

where *F*_1_*a*nd *F*_2_ are the production functions of each level, and τ _21_ *= τ*_2_/τ_1_ is the relative timescale of the two levels. This relative characteristic time gives an idea of how fast one level is related to the other. The periodic input γ(*t*) (see Methods for its description) stimulates a signaling component *y*_*1*_ (first level). In turn, *y*_*1*_ stimulates a second component *y*_*2*_ (second level). The behavior of each isolated level is known from the previous section and the second level’s activity does not directly affect the activity of the first one (cascade without feedback).

We use the notation *G*_1_ and *G*_2_ to represent the function *G*_*s*_ for the variables *y*_1_ and *y*_2_ respectively. To ensure dose conservation, we implemented amplitude compensation. We evaluated the frequency response across three relative time scales. Specifically, *y*_2_ was either slower (τ_21_ = 0.1), equal (τ_21_ = 1), or faster (τ_21_ = 10) than *y*_1_. The results are presented in Fig. 5.

**Figure 5.**
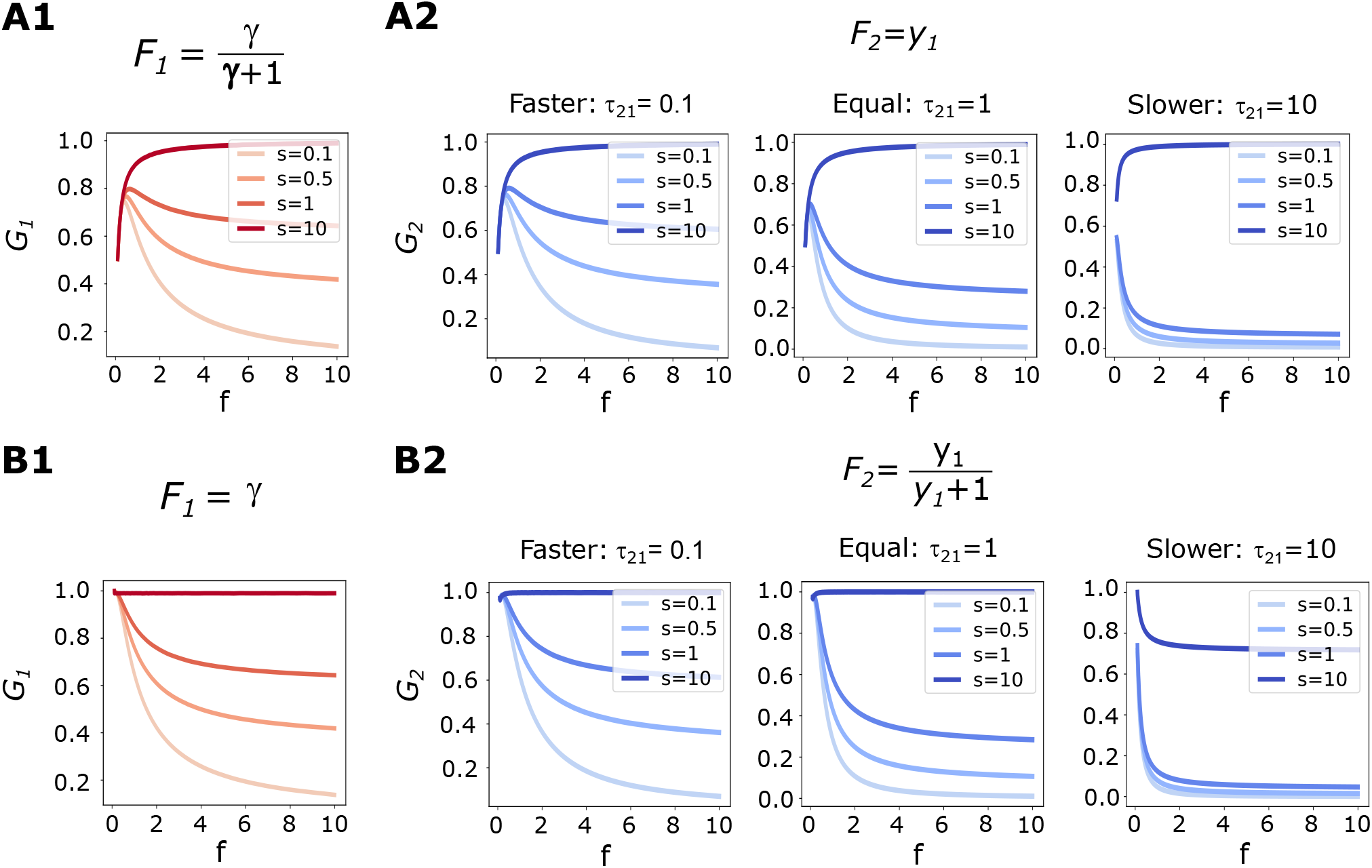
Transient frequency preference response is transmitted downstream in a feedforward signaling cascade depending on relative timescales. First level *y*_*1*_ is stimulated by a train of square pulses. Dose conservation is ensured by adapting the amplitude (reference input parameters: *d*_0_ = 0.1, *A*_0_ = 0.1, *T*_0_ = 1). The frequency response of each level was computed considering the observable *G*_*s*_, indicated as *G*_1_ or *G* _2_, respectively. The observable was normalized to its maximum value in the panel. Three scenarios were studied: a second level with a slower timescale than the first one (τ _21_= 0.1), *s*imilar timescales (τ_21_= 1), or faster (τ _21_ = 10). ***A***. Production function is concave for the first level, *F*_1_ *= γ*/(γ *+ 1*), and linear for the second one, *F*_2_*= y*_1_. Normalized *G*_1_ is plotted in red and normalized *G*^2^ in blue. Lines go from lighter to darker shades when *s* goes from earlier to later times. **B**. Frequency response for the two levels considering *F*_1_ *= γ, F*_2_ *= y*_1_/(*y*_1_ *+ 1*). The timescales and parameters are the ones used in panel A.

Initially, we analyzed the effects of a concave *F*_1_ function and a linear *F*^2^ function (Fig. 5A). As described in the preceding sections, *y*_1_ displays a transient frequency preference response. In isolation, *y*^2^ would function as a low-pass filter during both the transient and periodic regimes. However, different behaviors emerged in the cascade based on the relative timescale τ_21_. If τ_21_ *≤ 1, y*^2^ exhibited a transient band-pass filter and a high-pass filter in the periodic regime. Consequently, for relatively fast timescales, the response of the second level followed that of the first one, allowing the transmission of transient frequency preference downstream. Conversely, when the second level was slower than the first one τ_21_ *> 1*, the transient frequency preference was not transmitted effectively due to the dilution of the influence of the first level caused by the slower time scale. In Fig. 5B, *F*_1_ is linear and *F*_2_ concave. Results are analogous to the ones obtained in the previous study: the second level follows the frequency response of the first one, instead of behaving as if it was isolated.

Together, these results provide evidence that the frequency selectivity observed in the first level can be effectively transmitted to the downstream component in the feedforward cascade thereby altering the behavior the second level would have in isolation. However, when the second level operates at a slower pace relative to the first level, the former fails to reproduce the transient frequency preference. This finding supports the notion that a single level can convey two distinct messages downstream, depending on the dynamics of the subsequent level. Consequently, a second component characterized by a rapid response can decode the transient frequency preference, whereas a slower component is unable to do so. As a result, different interpretations of the same signal may arise in each scenario.

### 4. Ligand Receptor coupled to a Covalent Modification Cycle

In this section we consider a signaling network composed of a ligand receptor system coupled to a covalent modification cycle (CMC). This cycle is composed of two enzymatic reactions, working in opposite directions. In this cascade, the complex formed by the ligand-receptor acts as the kinase for the CMC. Fig. 6A shows a scheme of the system; the corresponding chemical reactions and ODE system is included in Methods. The receptor has four possible states: unbound (*R*), bound to the ligand (*RL*), bound to the substrate (*RS*), and bound to both (*RLS*). *RLS* is also the enzyme-substrate complex that catalyzes *S* activation into *S\**. The enzyme *E*_*d*_ forms a complex with *S*, ES\**, to deactivate it into *S*, completing the CM cycle (Di-Bella et al., 2018).

**Figure 6.**
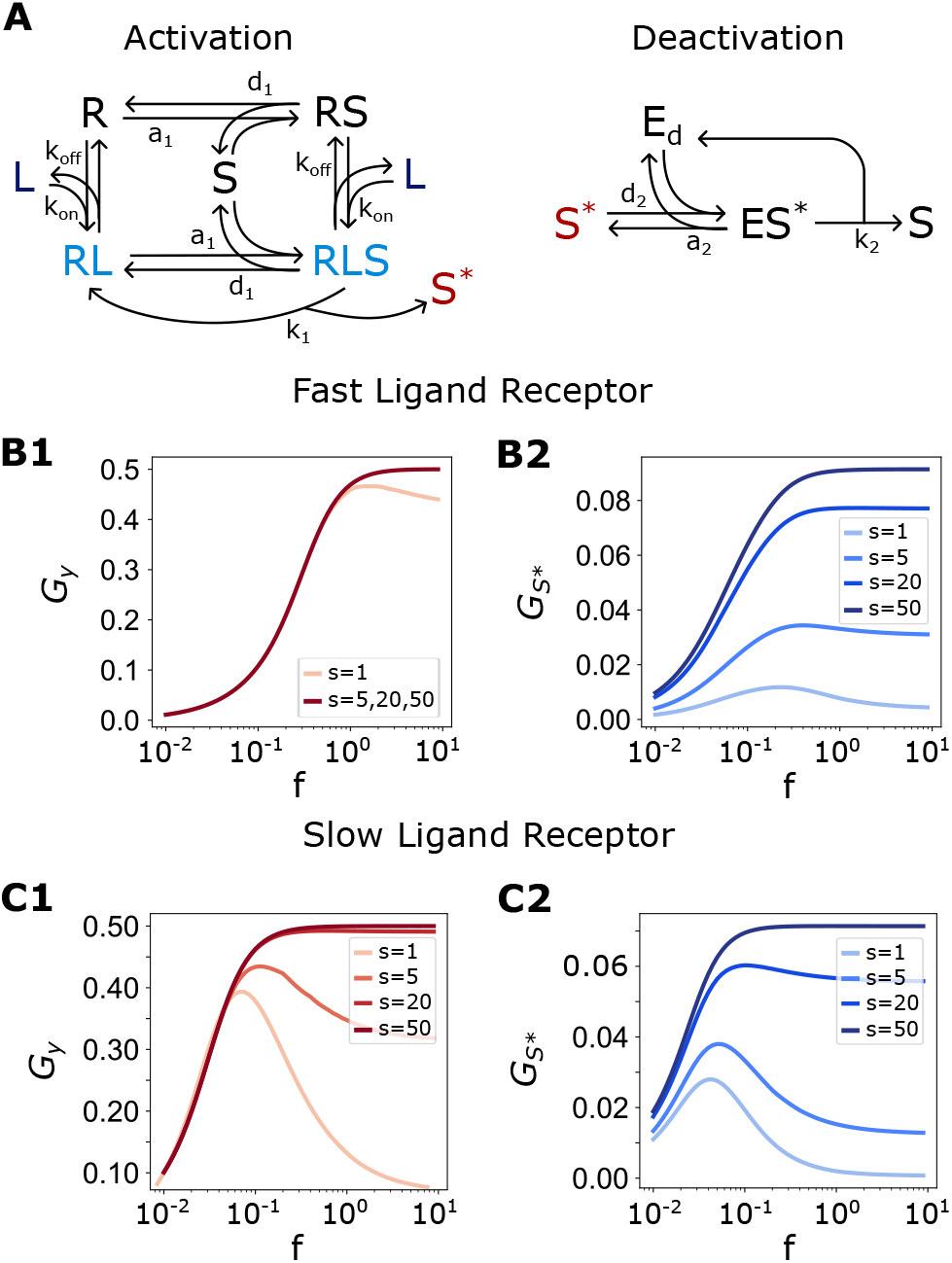
Frequency response in a feedfoward cascade with a ligand-receptor reaction coupled to a covalent modification cycle. **A**. Chemical reactions of the ligand-receptor system with a coupled covalent modification cycle. The stimulus is the ligand, and the output in the first level is the amount of ligand-receptor complex, given by *y = RL +RLS*. The output in the second level is the total amount of phosphorylated substrate *S*^*\**^. **B-C**. Frequency response for each level. The stimulation is a train of square pulses with dose conservation by amplitude (*d*_*0*_ = 0.*1, A*_*0*_ *=1, T*_*0*_= 1). The CMC parameters were kept constant: *k*_*1*_*=10, k*_*2*_*=10, a*_*1*_*=30, d*_*1*_*=30, a*_*2*_*=30, d*_*2*_*=300, S*_*T*_*=20, E*_*2T*_= 1. **B**. Observable *G*_*s*_ as a function of the frequency for the output *y* (**B1**) and for *S*^*\**^ (**B2**). Ligand-receptor parameters are *R*_*T*_ *=1, k*_*+*_*=1, k*_*-*_= 1. **C**. Observable *G*_*s*_ vs the stimulus frequency for *y* (**C1**) and for *S*^*\**^ (**C2**). Ligand-receptor parameters are *R*_*T*_*=1, k*_*+*_*=10, k*_*-*_*=0*.*1*, meaning that the reaction is slower than in panels B.

In this case, the periodic stimulus is the amount of free ligand of the system. The first level’s output is the total amount of the ligand-receptor complex, given by *y = RL + RLS*. The system is modeled in such a way that the CMC does not generate feedback over the ligand-receptor complex, thus the behaviour of this complex is not modified by the downstream reactions (demonstration in Supplementary Information, section 3). Then, the activity of the first level follows the characterization of the previous sections. The output of the second level is the phosphorylated substrate *S\**.

We explore how the frequency response in the first level is transmitted to the second one. We fixed the parameters related to the covalent modification cycle, while we varied the dimensional parameters of the ligand receptor system. For this study, CMC parameters were chosen so as the cycle response to a constant stimulus is monotonous (Di-Bella et al., 2018). We studied the frequency response of the CMC coupled to the ligand-receptor reaction in two scenarios: when the ligand-receptor reaction is slower than the isolated CMC and when it is faster. It is important to note that the time it takes the CMC to reach a steady state when it is coupled to a ligand-receptor system is higher than when isolated, as described in (Di-Bella et al., 2018). With the first set of parameters the timescale of the ligand receptor is slow compared to the CMC reactions. The timescale of the ligand receptor reaction is faster than the isolated CMC for the second parameter set, as the parameters values of the slow ligand-receptor were multiplied by 10. The ligand was stimulated with a train of square pulses considering dose conservation by amplitude compensation. The observable analyzed was the sliding window *G*_*s*_.

We compare the CMC frequency responses for different timescales in the upstream reaction. Results are shown in Fig. 6. Frequency responses in the scenario where the ligand-receptor is slower than the CMC are shown in panel B. Panel C exhibits the results for the case in which the ligand-receptor reaction is faster. It can be noted that the transient frequency preference of the fast ligand-receptor reaction (B1) is less pronounced than the one in the slow one (C1) for the same temporal window (determined by the parameter *s*). This is a consequence of the difference in the timescales: the ligand-receptor timescale also determines the rate at which the transient frequency response is lost. This behavior was not observed in the first section because a nondimensional system was considered, thus timescale effects were neglected.

In the panels corresponding to the CMC level (B2 and C2), we observe that the frequency responses are completely different even though the parameter set are identical. In Fig. 6-C2 the frequency preference is more persistent and pronounced than in panel B2. Although it takes the same time for the response amplitude to stabilize in both cases (between *s=20* and *s=50* the maximum value of the curve is modified), with similar characteristic times for the second signaling level, the curves have considerably different morphologies due to the behavior of the first reaction.

In panel B2, for *s > 5* the response in frequency is like a high-pass filter, with different maximum values as the time windows corresponds to later times. This is coherent with the results in panel B1, where from *s=5* the response of the ligand-receptor is a high-pass filter. On the contrary, in C2 there is a clear transient frequency response for *s=1* and *s=5*, which seems to last until *s=20*. This behavior is consistent with the response of the slow ligand-receptor system shown in C1, where the transient frequency response is more persistent and seems to be transmitted downstream.

From Fig. 6 we can conclude that the first level’s frequency response determines the response of the second node since changing the ligand-receptor timescales modifies the CMC frequency behavior.

### 5. MAPK cascade

This section focuses on the frequency response in a more intricate cellular signaling system known as the MAPK cascade. The model utilized in this study is based on the framework described in the work by Huang and Ferrell (Huang & Ferrell, 1996) and later adapted by Qiao et al. (Qiao et al., 2007). Illustrated in Fig. 7A, the model comprises three interconnected levels of covalent modification cycles. In the first level, a single phosphorylation occurs, while the second and third levels involve dual phosphorylation of the kinase. The phosphorylated substrate within each level, highlighted in color in Fig. 7A, functions as the kinase for the subsequent level. The model description is governed by mass action kinetics, resulting in fifteen differential equations and over thirty parameters. This system exhibits remarkable versatility, displaying a wide range of behaviors under constant stimulation, including bistability and oscillations, as demonstrated by Qiao et al (Qiao et al., 2007). In our study, we examined the frequency response of the system employing a parameter set reported in a previous study (Shankaran et al., 2009). The complete model and corresponding parameter values can be found in the Methods section. With this parameter set, the system’s response to a step-like stimulus is monotonous.

**Figure 7.**
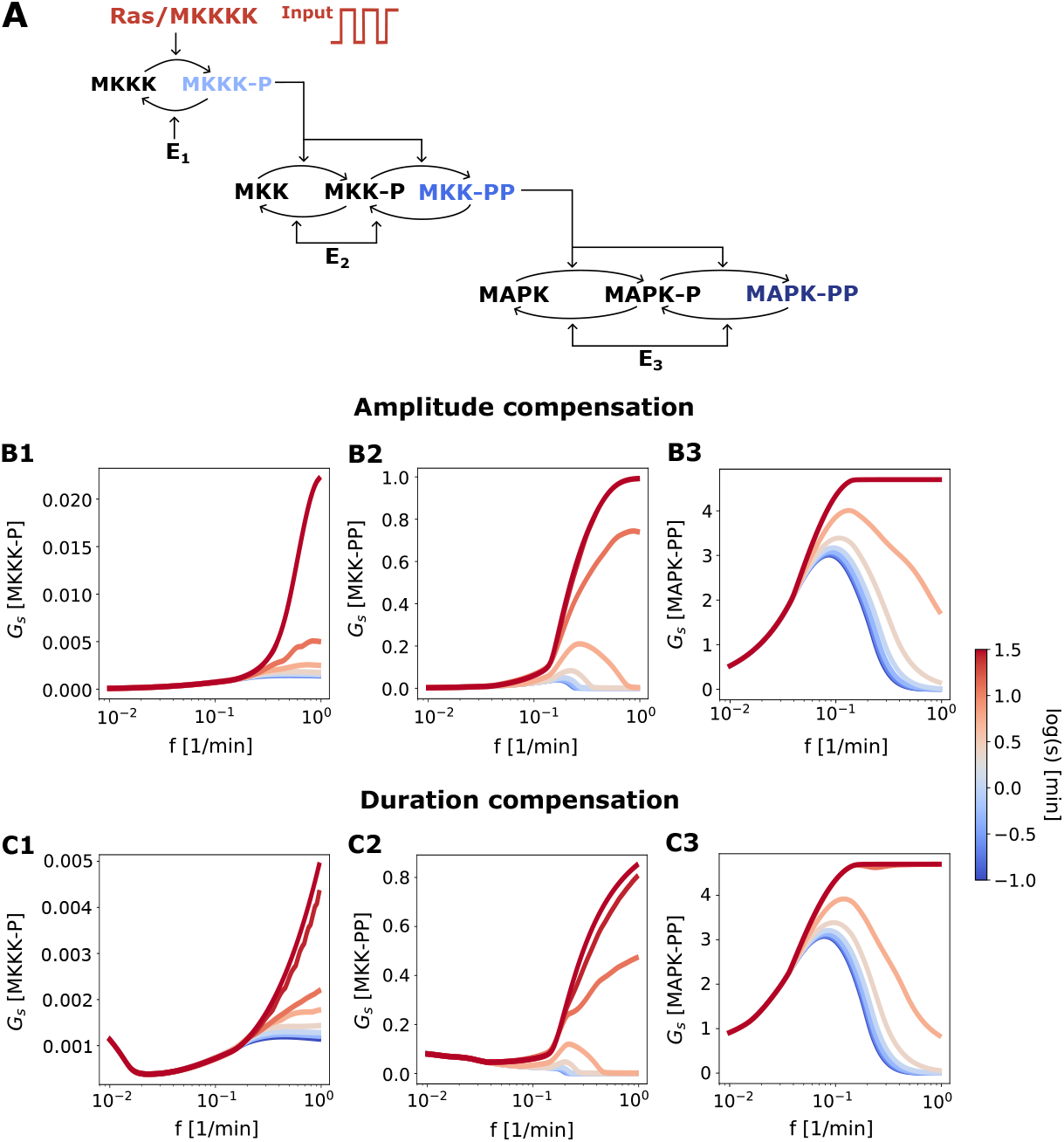
Transient frequency preference in a MAPK cascade. **A**. Scheme representing a three level MAPK cascade. The parameter set was taken from (Shankaran et al., 2009)(see Methods). A periodic stimulation is applied in the Ras/MKKKK level (with dose conservation by amplitude (panels B) and duration compensation (panels C). Input parameters: *A*_*0*_*=10, d*_*0*_*=1 min, T*_*0*_ *= 10 min*. The observable *G*_*s*_ is measured in the three levels MKKK-P, MKK-PP, MAPK-PP. The color of the curves indicates the initial time *s*, blue is related to transient time, whereas red indicates the periodic regime. Frequency response: **B1** and **C1** first level (MKKK-P), **B2** and **C2** second level (MKK-PP), **B3** and **C3** third level (MAPK-PP).

In this section, our focus is on analyzing the frequency response of the three levels within the signaling pathway. To achieve this, we employed a periodic input, representing the total amount of Ras/MKKKK in the system. To quantify the response, we measured the observable *G*_*s*_ for each of the three levels: MKKK-P, MKK-PP, and MAPK-PP. To stimulate the system, we utilized a train of square pulses with dose conservation through both amplitude and duration compensation.

When introducing variations in the total amount of a protein, it is important to consider mass conservation principles to ensure biological plausibility. In our study, we encountered situations where the calculated dose resulted in negative values, which are biologically nonsensical. To address this issue, we implemented an adaptation strategy for the stimulus, as in (Szemere et al., 2021). Specifically, at each time point, we stimulated the desired total amount of Ras/MKKKK and checked if the resulting dose was negative. If it was negative, we set the dose value to 0. This adaptation ensured that only meaningful and non-negative doses were considered in our simulations. Through extensive simulations, we observed that this adaptation did not significantly alter the overall total amount of Ras/MKKKK. The average total amount remained relatively constant across all periods examined. For further details and data, refer to Methods.

In this section, we investigate the frequency response of the MAPK cascade under the given conditions, and the findings are presented in Fig. 7. Similar results were obtained with both dose conservation protocols. As opposed to the first level (panels B1 and C1), both the second and third levels demonstrate a clear transient frequency response. Notably, the third level displays a selectivity in frequency during the transient phase, with a comparable width to the second level but exhibiting a higher amplitude relative to the steady-state response. For an analysis of the input parameter’s influence see Supplementary Information, section 6.

Thus, when considering the model of the MAPK pathway simulated with a parameter set derived from experimental data, we observe a transient frequency preference in the second level that becomes further amplified in the third level. As opposed to models considered in previous sections, this model includes hidden feedbacks (Ventura et al., 2008). Our results suggest that transient frequency preference may potentially be observed in more realistic models and that it can become more pronounced and enduring in downstream levels.

### 6. Frequency preference in a ligand-receptor system is also found without dose conservation

The transient nature of the frequency preference response in the ligand-receptor system, as observed through the sliding window *G*_*s*_ and the expanding window *H*_*s*_, under a dose conservation protocol, raises questions about its broader applicability. It remains unclear whether this phenomenon is exclusive to these specific characteristics or if it emerges under different conditions, such as in the periodic regime or in the absence of dose conservation. In this section, we study a different observable in the frequency response of a ligand-receptor system. Inspired by a previous experimental study (Bugaj et al., 2018), we focus on the integral gain over a threshold (*I*_*G*_) as the chosen metrics. By applying a periodic stimulation γ(*t*), we investigate the values of the complex *y* that surpass a specified threshold *u, y*_*threshold*_ (*t*) *= y*(*t*) *≥ u*, thereby defining the following observable:

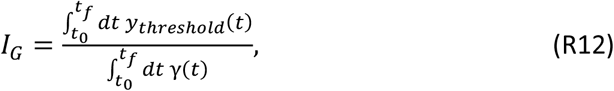

where *t*_*0*_ and *t*_*f*_ are the time integration limits. In this section, we depart from a dose conservation approach and, instead, calculate the integral gain by dividing by the integral of the stimulus. By computing the gain, valuable insights can be gleaned regarding the temporal dynamics and processing of signals within the ligand-receptor system, decoupled from changes in the overall input dosage.

Fig. 8 presents the frequency response of the dimensionless ligand-receptor system using the observable *I*_*G*_. The figure comprises panels displaying different combinations of integration time windows and thresholds.

**Figure 8.**
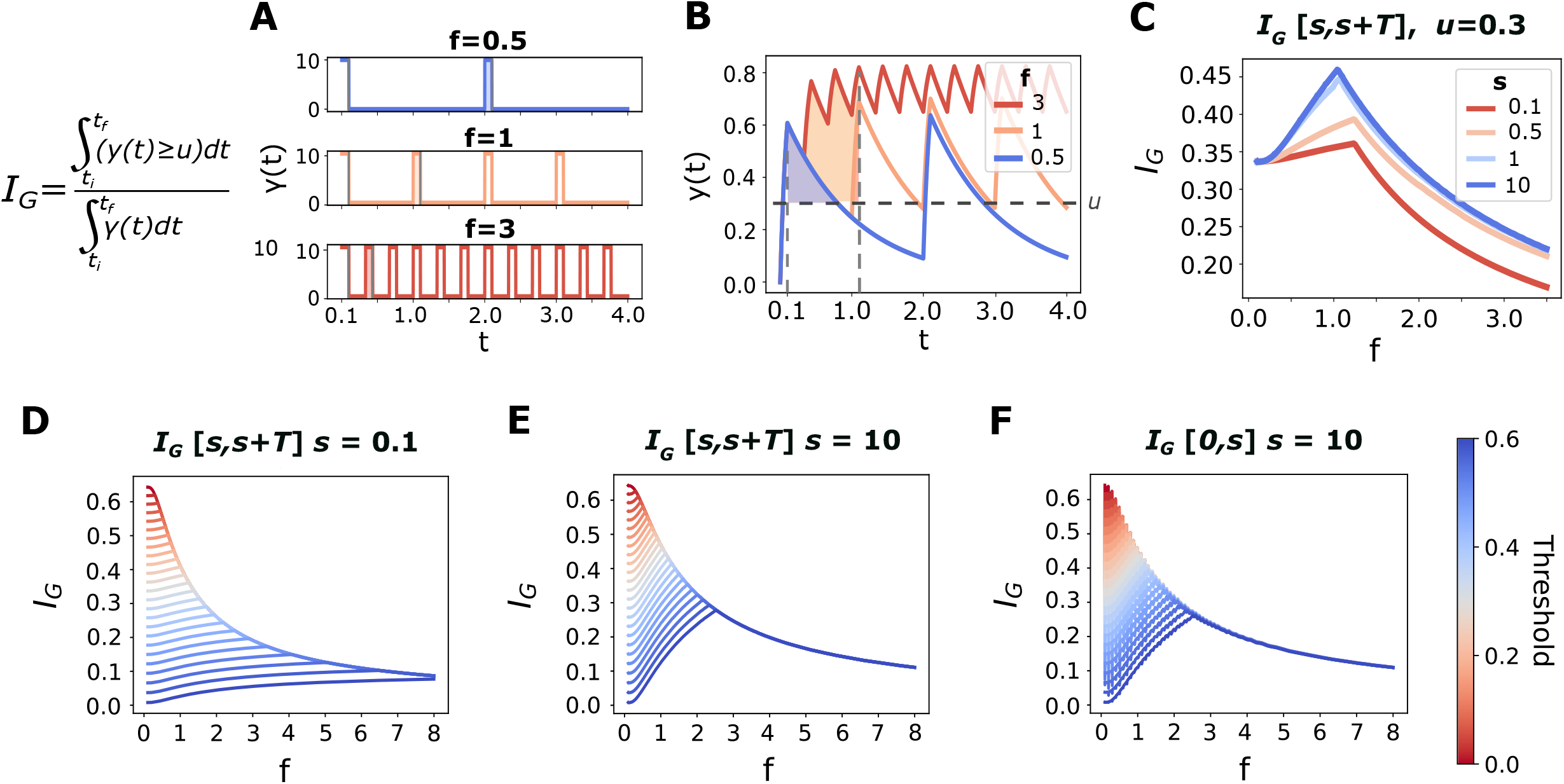
Frequency response for the ligand-receptor system for observable *I*_*G*_ . The observable *I*_*G*_ was calculated for different temporal windows by integrating the response above a threshold (see Eq. R12). Input parameters *A*_*0*_ *= 10, d*_*0*_*=0*.*1, T*_*0*_= 1. **A**. Input as a function of time for three different frequencies. **B**. The dimensionless ligand-receptor complex as a function of time for the three input frequencies in panel A. The shadowed area indicates the integral above a threshold (*u=0*.*3*) over a period from *s=0*.*1*. **C**. Frequency response for the observable *I*_*G*_ for different initial times. The value of the threshold is fixed in *u=0*.*3*. The observable was computed for a period-sized window for different initial times *s*, indicated by the color scale. **D. E. F**. Frequency response for the observable *I*_*G*_ for different temporal windows and thresholds. Panel D considers the observable *I*_*G*_ between *s* and *s+T*, for *s= 0*.*1*, corresponding to a transient response. Panel E displays the observable for a period sized window for *s=10*, corresponding to a pseudo-periodic regime. Panel F shows the frequency dependence for the observable *I*_*G*_ for an accumulated temporal window (from *0* to *s=10*).

Panels A and B indicate the input *γ*(*t*) and the output *y*(*t*) for three different frequencies. The shadowed area indicates the integral above a threshold over a period. In panels C, D and E, *I*_*G*_ was computed between *s* and *s+T*. Panel C explores the frequency response dependence of the observable *I*_*G*_ *a*s *s* is varied, for a fixed threshold. Panel D, E and F explore the frequency response dependence with the threshold value, and for each panel we consider a different temporal window. For panel F, *I*_*G*_ was calculated within the time window from *0* to *s*, with *s=10*, analogous to the observable *H*_*s*_. In panel D the temporal window goes from *s* to *s+T*, where *s=0*.*1* and *T* represents the stimulus period. At this value of *s*, the system exhibits a pseudo-periodic behavior. In panel E, *I*_*G*_ was evaluated between *s* and *s+T*, with *s=0*.*1*. At this value of *s*, the system exhibits a transient response. In these three panels, the color scale indicates different threshold values employed in the *I*_*G*_ calculation.

Frequency preferences only emerge when considering a nonzero threshold in the calculation of *I*_*G*_. When the observable is estimated without a threshold, the response is a low-pass filter for every time window (red color in panels D, E and F). Note that this behavior is different from results obtained in a dose conservation protocol. Moreover, increasing the threshold value generates more pronounced frequency preferences.

The ligand-receptor system exhibits a frequency preference response when the observable *I*_*G*_ is examined during both transient and periodic regimes when a threshold is imposed. This preference is more pronounced and of higher magnitude during the periodic regime. This behavior becomes evident when calculating *I*_*G*_ between *s* and *s+T*, as well as between 0 and *s*. In panel F slight variations are observed in certain curves. This phenomenon arises due to the calculation of *I*_*G*_ over incomplete periods, as previously explained for the observable *H*_*s*_ in Fig. 2. These small variations highlight the sensitivity of the ligand-receptor system to the precise duration of the time window considered when evaluating *I*_*G*_.

The results obtained in this section demonstrate a clear preference for specific frequencies in the ligand-receptor system. Remarkably, this preference arises without requiring dose conservation or restricting measurements to only the transient stage.

## Discussion

### Main ideas of the manuscript

In this study, our focus lies in investigating the frequency response of various systems that share a common characteristic: their response to a constant stimulus exhibits a monotonic behavior. Analyzing the frequency response of a system takes several factors into account. These factors include the choice of the observable being measured, the temporal window in which the measurement occurs, as well as the specific features of the stimulus itself, such as whether the dose is conserved or not and the mechanisms employed to achieve dose conservation.

Given the diverse range of factors at play, characterizing a system’s ability to exhibit frequency preferences can be a complex endeavor. Consequently, in this study, we opted to evaluate different simple systems, both in isolated and coupled configurations, under various parameter settings. Furthermore, we considered observables inspired by different experimental scenarios, thereby encompassing a broader spectrum of possibilities. This approach allows us to explore a multitude of scenarios and gain valuable insights into the behavior of these systems.

### Dose-conservation

In this paper, most of the presented results adhere to a dose conservation scheme, employing either amplitude or duration compensation, following the approach described in Fletcher et al. (Fletcher et al., 2014a). This method ensures that we are investigating genuine frequency encoding phenomena. By maintaining a consistent mean dose of stimulation across different frequencies, we mitigate the potential influence of variations in mean dose on observed differences in frequency response. This allows us to focus specifically on the decoding mechanism of temporal information rather than confounding effects stemming from changes in the mean dose.

### Observables

As mentioned previously, the choice of the measured observable depends on the specific biological context. In the literature, several observables have been identified as relevant in different studies. One such observable is the response amplitude, which naturally emerges in various contexts (Krivan et al., 2002b; Mettetal et al., 2008). It is defined as the difference between the maximum and minimum values of the response and provides valuable information about the magnitude of the cellular response. Another important observable is the gain, which represents the ratio of the stimulus amplitude to the response amplitude (Rotstein & Nadim, 2014; Vera et al., 2020). The gain quantifies the amplification or attenuation of the response relative to the stimulus and can provide insights into the system’s sensitivity. The integral of the response is also commonly considered (Wilson et al., 2017). It considers the accumulation of a substance that can trigger a signal, thus capturing the overall response over time. In certain cases, researchers may be interested in determining if the response exceeds a specific threshold. In such instances, the observable of interest becomes the integral of the response above a threshold (Bugaj et al., 2018), allowing for a focused analysis of events surpassing the designated threshold. Other studies have explored the average response (Fletcher et al., 2014; Hannanta-Anan & Chow, 2016), which involves normalizing the integral by the duration of the temporal window considered for the integration. The average response provides a measure of the system’s overall activity within the specified time frame. Overall, the selection of an appropriate observable depends on the specific research question and the desired aspect of the cellular response being investigated.

### Transient versus periodic regime

When investigating simple isolated systems stimulated with a train of square pulses that conserve dose, we observed the emergence of a phenomenon we called transient frequency response. This phenomenon manifests as a frequency selectivity that occurs when observables are measured before the system reaches its periodic regime. The transient frequency response holds biological significance as it may shed light on how a system can transmit distinct messages downstream based on the timing of the response measurement (Ventura et al., 2014). Furthermore, our findings also indicate that the phenomenon is not exclusive to persistent stimuli, as it remains of interest when the stimulus duration is relatively short compared to the intrinsic time of the stimulated system. This observation aligns with previous studies that have highlighted the importance of transient stimuli in biological systems (Blitz & Regehr, 2003; Marhl & Grubelnik, 2007; Raina et al., 2022). This suggests that the transient frequency response may play a significant role in various biological processes, warranting further exploration and investigation.

### Ligand-receptor system

We characterized the transient frequency response of a ligand-receptor reaction. The selectivity can be interpreted in two ways as the model is dimensionless: the time in our system is *t* = *t*′*k*_*off*_, then the frequency is actually *f = f*′/*k*_*off*_. *T*herefore, we can think that a given ligand-receptor system has an optimal response to an input, or that, given a fixed input we find a ligand-receptor system, characterized by its dissociation rate constant, that optimally responds to that input.

It should be noted that the ligand-receptor system, in addition to representing the chemical reaction of ligand-receptor, is a general model of reversible binding between two molecules (Ingalls, 2013). Moreover, it can also represent a simplified signaling model. For example, the ligand-receptor system is a limit case of covalent modification cycle, in the weakly activated regime (Heinrich et al., 2002; Szemere et al., 2021). Consequently, our findings for this model have implications that extend beyond the chemical reaction of a ligand-receptor.

### Simplified linear description

We studied the frequency response of a signaling component admitting a linear one-dimensional mathematical description as outlined in Fletcher et al. (Fletcher et al., 2014a). This model is also present in the literature as a feedfoward system as in Fletcher et al. or as simple regulation model as in several references (Alon, n.d.; Cournac & Sepulchre, 2009; Tran & Clayton, 2023). To ensure versatility in representing various biological scenarios, we examined different production rate functions denoted as *F*. Firstly, we considered a linear *F*, which corresponds to a linear synthesis term (Tchourine et al., 2014) and a weakly activated protein kinase cycle (Heinrich et al., 2002b). Secondly, a concave *F* emerged from Michaelis-Menten kinetics (Ma et al., 2009). Additionally, a sigmoidal *F* was considered, resulting from a Hill term (Cournac & Sepulchre, 2009). Furthermore, we encountered a convex *F* in the same scenario, but only for low inputs.

Based on our numerical and analytical findings (refer to Supplementary Material), it is evident that the frequency response is influenced by the type of nonlinearity present in the regulatory function. These results align with the observations reported by (Fletcher et al., 2014). Furthermore, we investigated the transient regime and discovered that concave and sigmoidal functions can lead to a frequency preference response, contingent upon the characteristics of the production function’s shape.

### Coupled signaling components without and with feedback

We conducted a study on the frequency response of simple systems integrated within complex signaling networks. Specifically, we examined two instances of a two-node cascade without feedback loops, ensuring that the activity of the second node does not influence the first node. In both cases, we discovered that the initial node’s transient frequency preference can be transmitted and subsequently determines the frequency preference of the following node. The transmission of the transient frequency response is contingent upon the relative timescales of the nodes. When the second node operates at a faster timescale than the first node, the transient frequency response is effectively conveyed.

We conducted an analysis to examine the emergence of transient frequency preference within a three-level MAPK-kinase cascade. The cascade was described mechanistically (Huang & Ferrell, 1996b) and the parameter values we used were experimentally reported (Shankaran et al., 2009). Our findings revealed the presence of frequency selectivity in both the second and third levels, with a notable amplification observed in the latter stage. Considering that temporal aspects play a crucial role in signal transduction within the ERK pathway (Alberts et al., 2019; Bugaj et al., 2018; Rauch et al., 2016; Ryu et al., 2015), the identification of transient frequency preference in the MAPK model holds potential relevance in the context of decision-making processes. Unlike the previous systems studied, the ERK pathway exhibits sequestration-induced implicit feedback between levels, making it an example of a network with retroactivity (Sepulchre et al., 2012; Ventura et al., 2008). This feedback mechanism likely contributes to the amplification of the transient frequency phenomenon.

### Ligand-receptor system with a different setting

Lastly, we investigated the frequency response of the ligand-receptor system under different hypotheses. We discovered the presence of frequency preference within the periodic regime, specifically when the observable considered is the integral above a threshold without dose conservation. This observation emphasizes the remarkable flexibility of a one-dimensional model, as it demonstrates the ligand-receptor system’s ability to exhibit frequency preference even in the periodic regime.

### Conclusions

The major contribution of this work lies in the study of frequency responses considering a temporal window during which the output of the stimulated system is in a transient phase. To the best of our knowledge, there are no existing theoretical precedents employing a similar approach, except for the articles by Mahrl et al. (Marhl et al., 2005, 2006). However, it should be noted that their approach mixes transient and periodic regimes.

Our findings provide support for the hypothesis that frequency preference is not an inherent property of a system and the connectivity between nodes, but instead emerges from the dynamic interplay between the stimulus and response, how the response is defined and when it is measured. This highlights the remarkable versatility of biological systems, enabling the transmission of multiple messages through a single signal while also exhibiting high specificity based on the characteristics of the signal-processing system.

## Methods

List and definitions of attributes analyzed in the article.

**γ** *i*nput

**T** input period

**f** input frequency

**d** pulse duration

**A**_**0**_ pulse amplitude corresponding to an input of a single pulse (reference input)

**d**_**0**_ reference pulse duration

**T**_**0**_ reference period

**y** output

**τ** *t*ime to reach steady state under constant stimulation

**τ**_**p**_ *t*ime to reach a periodic regime under periodic stimulation

**G**_***s***_ *s*liding window: temporal average of the output in a time interval that lasts one period of the input starting at time *s*, thus the time window is [*s, s*+T]

***H***_***s***_ *e*xpanding window: normalized accumulated integral up to a time *s*

***f***_***G***,***max***_ frequency leading to a maximum in the *G*_*s*_versus frequency curve (preferred frequency)

***G***_***max***_ maximum in the G_S_ versus frequency curve

**τ**_21_ *t*imescale associated to *y*_2_ relative to the timescale associated to *y*_1_

### Model equations

#### Ligand receptor system

The system equation, considering that the total amount of receptor is constant, and that the concentration of free ligand is not depleted significantly by the binding reaction is given by:

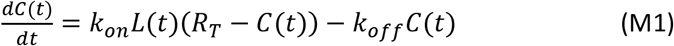

The decay time of the signal can be estimated when the stimulus is off: *L* = 0, thus: 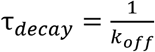. Nondimensionalized ODE equation for the complex *y*(*t*’), where *γ*(*t*’) is the scaled amout of ligand being pulsed is given by:

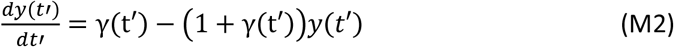

All the analytical calculations are performed over the dimensionless equation. As the variable and time are expressed in terms of a reference value, *R*_*T*_ for the complex concentration and *τ*_*decay*_ for the time, then in the frequency analysis the frequency is also scaled with the reference time. To simplify notation, dimensionless time *t’* is noted as *t* for the ligand-receptor analysis.

#### Signaling component with a simplified linear description

Dimensionless concentration of a given molecule *y*, with a production described by a function *F* and linear degradation. The system is stimulated with an input γ(*t*).

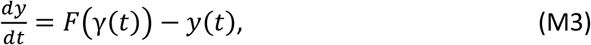

#### Coupled signaling components without feedback connections

Feedforward coupling, where a periodic input γ(*t*) stimulates a signaling component *y*_*1*_. At the same time, *y*_*1*_ stimulates a second component *y*_*2*_.

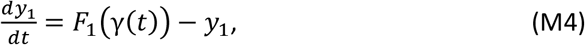

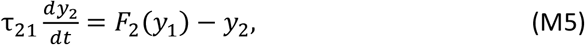

where *F*_*1*_ and *F*_*2*_ are the production functions of each level, and τ_21_ *= τ*^2^/τ_1_ is the relative timescale of the two levels. This relative characteristic time gives an idea of how fast one level is related to the other.

#### Ligand Receptor coupled to a Covalent Modification Cycle

The full set of reactions for this model is composed of those for a ligand receptor (LR) system and those for a CMC, with an extra reaction involving the binding of the receptor and the substrate without the ligand (Di-Bella et al., 2018b)

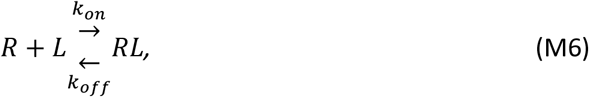

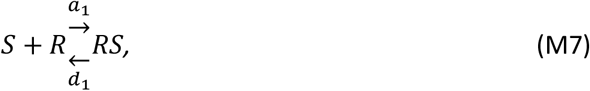

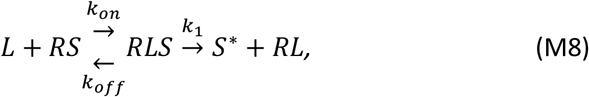

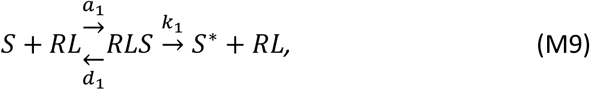

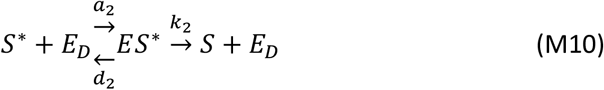

According to mass action kinetics, the ODE system describing the network is given by:

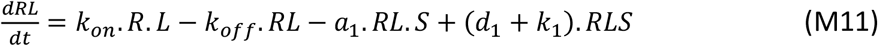

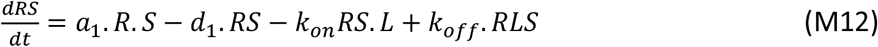

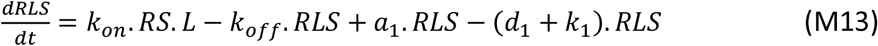

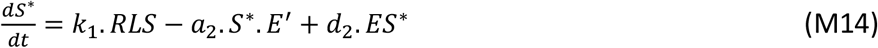

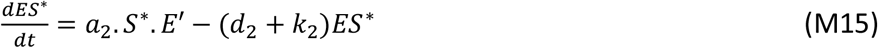

While the conservation laws are:

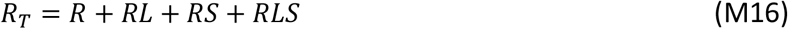

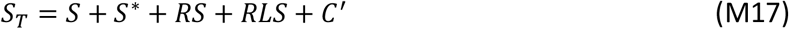

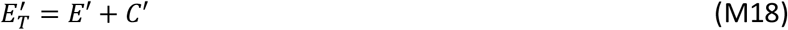

#### MAPK cascade

In the three levels MAPK cascade the first kinase *D*_*1*_ is periodically stimulated. Then, the cascade is composed of phosphatases E_1_, E_2_, E_3_; complexes C_1_-C_10_; MKKK molecule, noted as A and A* when phosphorylated (MKKK-P), the molecule MKK, M_0_ in ground state, M_1_ when one phosphate group is added (MKK-P) and M_2_ when two groups are added (MKK-PP); the molecule MAPK is noted as N_0_ in ground state, N_1_ when one phosphate group is bounded (MAPK-P) and N_2_ when two are added (MAPK-PP). The system equations are:

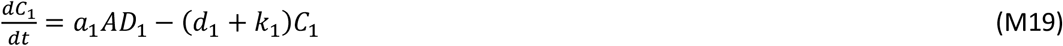

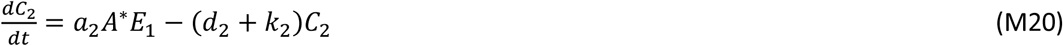

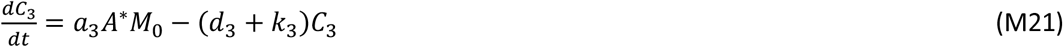

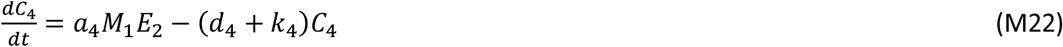

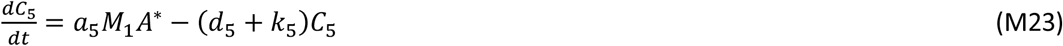

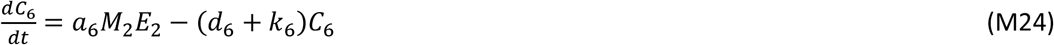

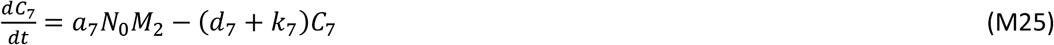

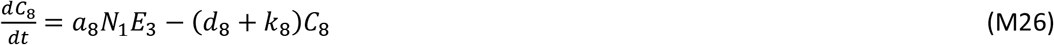

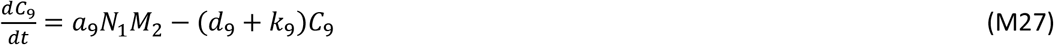

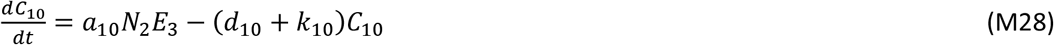

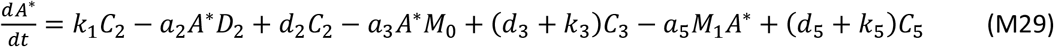

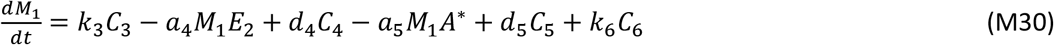

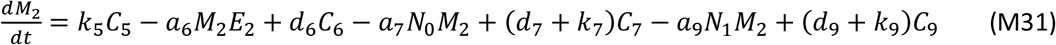

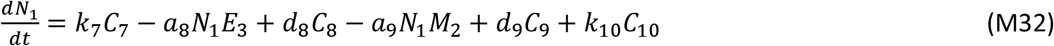

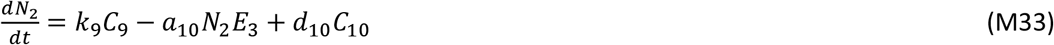

Mass conservation laws are given by:

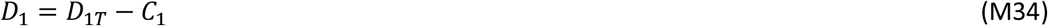

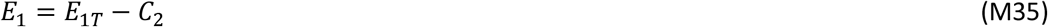

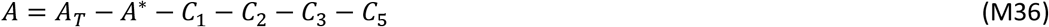

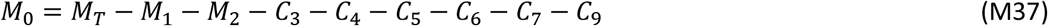

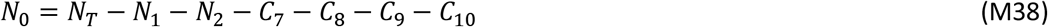

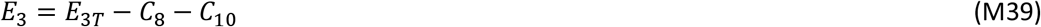

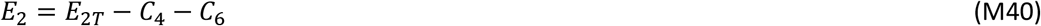

The parameters for simulations were taken from Shankaran et al. (Shankaran et al., 2009) given by: a_1_ = 1500 *μM, d*_1_ *= 4*.5 */min, k*_1_ = 10.5 */min* ; a_2_ = 1500 *μM, d*_2_= 1.5 */min, k*_2_ = 10.5 */ min* ; a_3_ = 1500 *μM, d*_3_ *=* 12 */min, k*_3_ = 10.5 */min* ; a_4_ = 1500 *μM, d*_4_ = 1^2^ */min, k*_4_ *=* 10.5 */min* ; a _5_ = 1500 *μM, d*_5_ *=* 12 */min, k*_5_ = 10.5 */min* ; a_6_ = 1500 *μM, d*_6_ = 12*/min, k*_6_ = 10.5 */min* ; a_7_ = 1500 *μM, d*_7_ *=* 12 */min, k*_7_ = 10.5 */min* ; a_8_ = 1500 *μM, d*_8_ *=* 12 */ min, k*_8_ = 10.5 */min* ; a_9_ = 1500 *μM, d*_9_ *=* 12 */min, k*_9_ = 10.5 */min* ; a_10_ = 1500 *μM, d*_10_ *=* 12 */min, k*_10_ = 10.5 */min* ; *A*_*T*_ = 0.1 *μM, D*_1*T*_ = 0.2 *μM, M*_*T*_ *=* 1.2*μM, N*_*T*_ *= 4*.8 *μM, E*_1*T*_ *=* 0.012 *μM, E*_2*T*_ = 0.06 *μM, E*_3*T*_= 0.05 *μM*.

### Timescales and parameters

We consider systems that are not autonomous oscillators (except for the case of the MAPK cascade, that can behave as an oscillator for particular parameter values). These systems are characterized by a timescale τ, which is the time to reach steady-state when stimulated with a step-like input, and are simple enough to produce, when forced with a periodic input, an output that can reach a pseudo-periodic regime. See formal proof of this last statement for simple systems in the Supplementary Information, section 1. We define τ_*p*_ as the time it takes such a system to reach the psedo-periodic solution. As proven in the Supplementary Information for simple systems, τ_*p*_ is a function of the period of the input, T. Since τ_*p*_ *c*an be bounded for every choice of T *c*onsidered in this article (see Supplementary Information, section 1 for more details), τ is a good estimator of the transient regime of the forced output.

The parameters involved in the first models (ligand-receptor, signaling components with a simplified linear description, and feedfoward cascade) are dimensionless. For example, in the ligand receptor system, a unit of time *t’* corresponds to 1/*k*_*off*_. Therefore, the frequency *f* is also scaled by the reference time, so that a unit of frequency corresponds to *k*_*off*_. *T*he concentration of the input is scaled by the dissociation constant for the binding-unbinding process and the complex is scaled by the total amount of receptor (R_*T*_).

For the rest of the explored models, some parameters have dimensions of concentrations (for example *R*_*T*_, *D*_*1T*_, *A*_*T*_), some have dimensions of 1/time (*k*_*off*_, *k*_1_) and others have dimensions of 1/(time _x_ concentration) such as *k*_*on*_ or *a*_1_. The unit of time can be selected as seconds or minutes so that the responses are consistent with different biological processes, and the unit of concentration is arbitrary. This means that when a value for a parameter is listed, the corresponding unit has to be added, for example, if the value of parameter a is 10, it has to be read as 10 × 1/(min _×_ concentration). Once the reference concentration is selected, all the parameters follow that reference unit. The interpretation of the results depends on the choice of the reference unit concentration, as the choice of the reference unit concentration remains a degree of freedom in our numerical methodology. For the MAPK cascade, parameters units are fixed, given by (Shankaran et al., 2009).

### Periodic input with dose conservation

For each model considered in this article we have a single ODE or a system of coupled ODEs describing the temporal evolution of the variables of interest. Let’s assume that the variable *y* is the output. The ODE/ODE system receives an input γ(t) *t*hat is a train of square pulses, defined as follows:

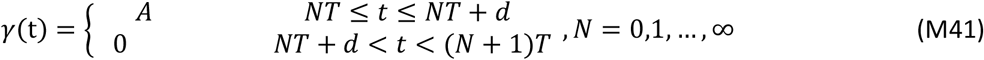

where *A* is the amplitude of the pulse, *T* is the period, *d* is the duration of the pulse and **N** is an integer number that indicates the number of the pulses.

We study how the system responds to a periodic input, focusing on its dependence on the frequency of stimulation. In our approach, similar to the one used in (Fletcher et al., 2014) the output is studied for different input frequencies in such a way that the average input per period is kept constant for all the frequencies. This approach is known as **input dose conservation**.

Dose conservation is accomplished through two protocols, depending which parameter is modified to keep constant the mean input as frequency is varied: by amplitude or by duration compensation. Both protocols start from a reference stimulus with an amplitude *A*_0_, pulse duration *d*_0_ and period *T*_0_. The mean dose *D* in the *N*^th^ period is:

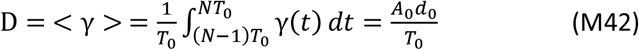

#### Dose conservation by amplitude compensation

Dose conservation is guaranteed by adapting the stimulus amplitude between one period an another, while fixing the duration of the on-phase of the pulse - *d*_*0*_ -, hence making the stimulus average to be constant for all the frequencies.

If we consider a new stimulus with a period T, to keep the mean dose constant, the amplitude of this input should be:

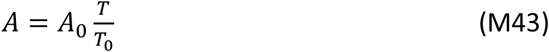

If we compute the mean dose in the N^th^ period, we obtain:

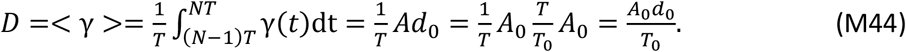

Thus, by this protocol, mean dose over a period is conserved as the frequency is varied.

#### Dose conservation by duration compensation

In this protocol, dose conservation is achieved by adapting the duration of the pulse, with a fixed amplitude *A*_*0*_, as the frequency is varied. Thus, for a period *T*, the duration must be:

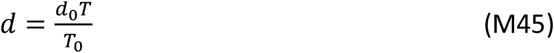

Then, the mean dose in a period results:

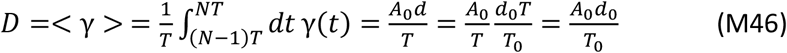

Note that in both protocols, the mean dose is constant if it is computed over an integer number of periods.

#### Adapted periodic input for the MAPK cascade

For the MAPK cascade system, the periodic input is the total amount of Ras/MKKKK (*D*_*1*_ in the model). In a closed system, the chemical variables are subject to mass conservation laws (Ingalls, 2014). The total amount of a substance must consider all the species where that substance participates in all the reactions that describe the system. For example, if this substance *X* has two possible states, *X* and *X’*, and it could also bind to another molecule forming a complex *C*_*X*_, then the total amount of this substance is *X*_*T*_ *= X + X*’ *+ C*_*X*_, where *X, X’* and *C*_*X*_ are variables in the ODE system. The presence of conservation laws conditions the periodic stimulation, since we introduce variations in the total amount of a protein, it is possible that the dose turns out negative, which has no biological sense. To vary the total amount *X*_T_, we adjust the value of the variable *X* for each time in the ODE system in a way such that according to the conservation law, the stimulus is the one desired. As *X’* and *C*_*X*_ depends on the systems dynamics, for example if we start from a positive value of *X*_*T*_ and the initial conditions are such that *X’* takes a high value, it is impossible to fix *X*_*T*_*=0* instantly, as in a train of square pulses, since *X* should be negative to compensate *X’* and *C*_*X*_. Therefore, adapted (modified) versions of the input must be used (Szemere et al., 2021).

The stimulus was adapted by stimulating the desired total amount of Ras/MKKKK at each time and setting its value to 0 if it resulted negative. From simulations we concluded that this adaptation does not severely modify the total amount of Ras/MKKKK, as its average is mostly constant in all periods considered.

### Observables for the frequency analysis

In this article, we focus on the analysis of the systems response to frequency, by means of different operations over the output.

#### Temporal averages

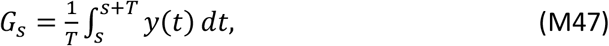

and

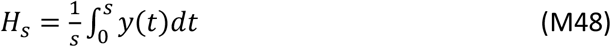

where *y(t)* is the temporal course of the output, *s* in an arbitrary initial time, T the stimulus period. For each fixed value of *s*, G_*s*_(*T*) is then a function of *T t*hat measures the temporal average of the output in a time interval that lasts for one period of the input starting at *s*, while H_*s*_(*T*) is the normalized accumulated integral. We call them sliding window and expanding window, respectively. G_*s*_(*T*) *c*an be computed during the transient part of the output, by choosing *s < τ*_*p*_, or when *y* has reached a periodic regime, by choosing *s > τ*_*p*_ (see Fig. 1). As mentioned above, τ is a good estimator of τ_*p*_ and it is easier to compute.

#### Integral gain over a threshold

We investigate the values of the complex *y t*hat surpass a specified threshold *y*_*threshold*_(*t*), thereby defining the following observable:

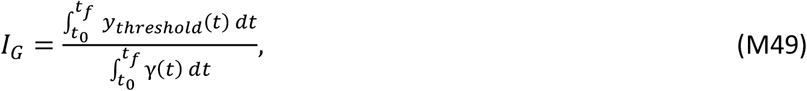

where *γ*(*t*) represents the stimulus, *t*_*0*_ and *t*_*f*_ the integration times.

## Acknowledgements

This work was supported by a grant from the from the Argentine Agency of Research and Technology (PICT2019-01681) to ACV and CSV. We thank Horacio Rotstein and Carla Bosia for fruitful discussions.

## Author contributions

ACV designed the project. CLS and JRS performed mathematical analysis and numerical simulations. RB and CSV performed analytical calculations. All the authors analyzed the results. CLS and ACV wrote the manuscript.

