## Supplementary Material for "Transient frequency preference responses in cell signaling systems"

### SUPPLEMENTARY INFORMATION

Contents


1. Mathematical framework 1
1.1. Ligand receptor model 2
1.2. Signaling components with a simplified linear description 4
1.3. Coupled signaling components without feedback connections 5
2. The signaling component with a simplified linear description response to frequency depends on the concavity of the function $F$ 6
3. Coupled covalent modification cycle does not generate feedback over the ligand-receptor system 11
4. Additional results for the ligand-receptor model 12
4.1. Transient frequency preference is also observed for a different periodic stimulation 12
4.2. Dose dependence in frequency response in the ligand-receptor system 12
5. Additional results for signaling components with a simplified linear description 14
5.1. Expanding window results 14
6. Additional results for the MAPK cascade model 14
References 17


#### 1. Mathematical framework

For each model considered in this article we have a single ODE or a system of ODEs describing the temporal evolution of the variables of interest. Lets assume that the variable $y$ is the output. The ODE/ODE system receives an input $\gamma$ that is a train of square pulses with period $T$. We define $\gamma$ a $T$ periodic function as follows:

$$\gamma(t) = \begin{cases} A_0T & if \quad t \in [0, d_0] \\ 0 & if \quad t \in (d_0, T] \end{cases}, \tag{1.1}$$

In this section we will analyze the time scales and asymptotic behavior of the following three models considered in the article:

**Ligand-receptor:**

$$\frac{dy}{dt}(t) = \gamma(t) - (1 + \gamma(t))y(t) \tag{1.2}$$

**Signaling components with a simplified linear description:**

$$y'(t) = F(\gamma(t)) - y(t). \tag{1.3}$$

**Coupled signaling components without feedback connections:**

$$y_1'(t) = F_1(\gamma(t)) - y_1 \tag{1.4a}$$

$$\tau_{21}y_2'(t) = F_2(y_1) - y_2 \tag{1.4b}$$

We consider only systems that are not autonomous oscillators. The systems we consider are characterized by a time-scale $\tau$ (the time to reach steady state when stimulated with a step-like increase of input) and are simple enough to produce, when forced with a periodic input, a periodic output of the same period and in phase with the input. We define $\tau_p$ as the time it takes such a system to reach the periodic solution.

In this section we will prove that the considered systems when forced with a $T$-periodic intput produce an asymptotically $T$-periodic output. Also, for each system we will compute the time $\tau$ it takes the system to reach the steady state when stimulated with a constant input conserving dose $\gamma(t) \equiv A_0 d_0$ and the time $\tau_p$ it takes the system to *reach the periodic solution* when forced with the step $T$-periodic function $\gamma$. This last time $\tau_p$ is clearly dependent on the period $T$. It is expected that for $T$ big enough, that is for low frequencies, the time $\tau_p$ will be large and for $T$ small, that is for high frequencies, the time $\tau_p$ will be small.

We first recall a known result for nonhomogeneous linear ordinary differential equation.

**Lemma 1.1.** *Given $f, g : \mathbb{R} \to \mathbb{R}$ continuous functions, then all the solutions of the*

$$z'(t) = g(t)z(t) + f(t) \tag{1.5}$$

*are given by*

$$z(t) = e^{\int_0^t g(s)ds} \left( \int_0^t f(s) e^{-\int_0^s g(r)dr} ds + k \right), \text{ with } k \in \mathbb{R} \text{ a constant.} \tag{1.6}$$

*If we have an initial datum $z(0) = z_0$, then the unique solution satisfying equation (1.5) and the initial datum is*

$$z(t) = e^{\int_0^t g(s)ds} \left( \int_0^t f(s) e^{-\int_0^s g(r)dr} ds + z_0 \right). \tag{1.7}$$

*Proof.* This is a classic result for linear ordinary differential equations. □

**Corollary 1.1.** *Given $f, g : \mathbb{R} \to \mathbb{R}$ continuous and $T$-periodic functions. If $z$ is a solution of the linear ordinary differential equation (1.5), then $u(t) = z(t + T) - z(t)$ satisfies the equation*

$$u'(t) = g(t)u(t),$$

*and therefore all the solutions are given by*

$$u(t) = k e^{\int_0^t g(s)ds}, \text{ with } k \in \mathbb{R} \text{ a constant.} \tag{1.8}$$

*Moreover, if $z(0) = 0$, the solution is*

$$u(t) = z(T) e^{\int_0^t g(s)ds}. \tag{1.9}$$

*Proof.* Note that if $z$ is a solution of (1.5), from the periodicity of $f$ and $g$, then

$$z'(t + T) = g(t)z(t + T) + f(t), \tag{1.10}$$

and therefore substracting equations (1.5) and (1.10)

$$\frac{d}{dt}(z(t + T) - z(t)) = g(t)(z(t + T) - z(t)). \tag{1.11}$$

From lemma 1.1 and using that $u(0) = z(T) - z(0)$, we obtain the remaining of the proof. □

#### 1.1. **Ligand receptor model.**

##### 1.1.1. **The solution is asymptotically periodic.** 

Following we prove that the solution of the ligand receptor model (1.2) is an asymptotically periodic function.

**Proposition 1.1.** *Given $\gamma(t)$ a $T$-periodic function. Then the solution of equation (1.2) is an asymptotically $T$-periodic function.*

*Proof.* Since the solution satisfies equation (1.2) and $\gamma$ is a $T$-periodic function, from corollary 1.1 we deduce that

$$\frac{d}{dt}(y(t + T) - y(t)) = -(1 + \gamma(t))(y(t + T) - y(t))$$

and there exists a constant $k \in \mathbb{R}$ such that

$$y(t + T) - y(t) = k e^{-\int_0^t 1+\gamma(r)dr}$$

which goes to 0 when $t \to \infty$ since $\gamma$ is bounded. □

**Remark 1.1.** In particular, for a periodic pulse as in (1.1) and taking initial datum $y(0) = 0$, since the solution of (1.2) is given by

$$y(t) = \int_0^t \gamma(s) e^{-\int_s^t 1+\gamma(r)dr} ds \tag{1.12}$$

we have that

$$y(T) = \int_0^T \gamma(s) e^{-\int_s^T 1+\gamma(r)dr} ds = \frac{A_0 T}{1+A_0 T} e^{-T}(e^{d_0} - e^{-A_0 T d_0}).$$

Thus, from (1.9)

$$y(t+T) - y(t) = y(T) e^{-\int_0^t 1+\gamma(r)dr} = \frac{A_0 T}{1+A_0 T} e^{-T}(e^{d_0} - e^{-A_0 T d_0}) e^{-t} e^{-\int_0^t \gamma(r)dr}. \tag{1.13}$$

1.1.2. **Time scales.** We now analyze the time $\tau$ for the LR model. When the system is forced with a constant input $\gamma(t) \equiv A_0 d_0$, the solution (1.12) becomes

$$y(t) = A_0 d_0 \int_0^t e^{-(1+A_0 d_0)(t-s)} ds = \frac{A_0 d_0}{1+A_0 d_0}\left(1 - e^{-(1+A_0 d_0)t}\right).$$

and therefore the system takes time $\tau = \frac{1}{1+A_0 d_0}$ to reach $(1 - e^{-1})$ times the steady state $\frac{A_0 d_0}{1+A_0 d_0}$.

If we use the approximation

$$\int_0^t \gamma(r)dr \sim A_0 d_0 t,$$

we deduce from (1.13)

$$y(t+T) - y(t) \sim \frac{A_0 T}{1+A_0 T} e^{-t(1+A_0 d_0)} e^{-T}\left(e^{d_0} - e^{-A_0 T d_0}\right). \tag{1.14}$$

Now we can estimate the time $\tau_p$. We take $\varepsilon > 0$ small enough such that $y(0+T) - y(0) = y(T) > \varepsilon$. Using (1.14) for $t = 0$, we choose $\varepsilon$ such that

$$\frac{A_0 T}{1+A_0 T} e^{-T}\left(e^{d_0} - e^{-A_0 T d_0}\right) > \varepsilon \tag{1.15}$$

Then, the time $\tau_p$ that it takes to sastisfy $|y(t+T) - y(t)| < \varepsilon$ for all $t > \tau_p$ is given by

$$\tau_p(T) \sim \frac{\ln\left(\left(e^{d_0} - e^{-A_0 T d_0}\right) e^{-T} \frac{A_0 T}{1+A_0 T} \varepsilon^{-1}\right)}{1+A_0 d_0} = \ln\left(\left(e^{d_0} - e^{-A_0 T d_0}\right) e^{-T} \frac{A_0 T}{1+A_0 T} \varepsilon^{-1}\right)\tau \tag{1.16}$$

Note that for each $T$ such that (1.15) is satisfied we have that $\tau_p > 0$. Moreover, since $\frac{A_0 T}{1+A_0 T} \le 1$ we obtain

$$0 < \left(e^{d_0} - e^{-A_0 T d_0}\right) e^{-T} \frac{A_0 T}{1+A_0 T} \le e^{d_0} e^{-T} \le 1. \tag{1.17}$$

Then,

$$\tau_p(T) < \frac{\ln(\varepsilon^{-1})}{1+A_0 d_0} = \frac{-\ln(\varepsilon)}{1+A_0 d_0} = \tau_{p,max}.$$

Thus, if we assume that the constants $A_0, d_0$ satisfy $\varepsilon^{-1} \le e$ $(\varepsilon > e^{-1})$ then

$$\tau_p(T) \le \frac{1}{1+A_0 d_0} = \tau. \tag{1.18}$$

Moreover, for $\varepsilon > 1$ we have that $\tau_p < 0$ and also for all $\varepsilon > 0$, $\tau_p(T) \to -\infty$ when $T \to \infty$.

1.1.3. $\tau_{p,max}$ **for** $T$ **bounded.** Conversely, if $T$ **is bounded**, for instance assume $d_0 < T < T_{max}$. Using that $e^{-T}$ is decreasing and no negative and $\left(e^{d_0} - e^{-A_0 T d_0}\right)$, $\frac{A_0 T}{1+A_0 T}$ are both no negative and increasing functions with respect to $T \in [d_0, T_{max}]$ then

$$\left(e^{d_0} - e^{-A_0 d_0^2}\right) e^{-T_{max}} \frac{A_0 d_0}{1 + A_0 d_0} < \left(e^{d_0} - e^{-A_0 T d_0}\right) e^{-T} \frac{A_0 T}{1 + A_0 T} \tag{1.19}$$

Then if we assume that the constants $A_0, d_0, T_{max}$ satisfy

$$\varepsilon < \left(e^{d_0} - e^{-A_0 d_0^2}\right) e^{-T_{max}} \frac{A_0 d_0}{1 + A_0 d_0}$$

we have that $\tau_p(T) > 0$ for all $T$. Also, if we assume that

$$\varepsilon < \left(e^{d_0} - e^{-A_0 d_0^2}\right) e^{-T_{max}} \frac{A_0 d_0}{1 + A_0 d_0} e^{-1} \tag{1.20}$$

we deduce from (1.16) and (1.19) that $\tau_p(T) > \tau$ for all $T \in [d_0, T_{max}]$.

1.2. **Signaling components with a simplified linear description.** For this model we assume that the function $F$ is derivable, positive in $(0, +\infty)$ and $F(0) = 0$. Solving the ode (1.3), we obtain

$$y(t) = \int_0^t F(\gamma(r)) e^{-t+r} dr, \tag{1.21}$$

1.2.1. **The solution is asymptotically periodic.** Following we prove that the solution of the simple synthesis and degradation model (1.3) is an asymptotically periodic function.

**Proposition 1.2.** *Given $F : \mathbb{R} \to \mathbb{R}$ a continuous function and $\gamma(t)$ a $T$-periodic function. Then the solution of equation* (1.3) *is an asymptotically $T$-periodic function.*

*Proof.* Since $y$ satisfies equation (1.3) and $\gamma$ is a $T$-periodic function, from corollary 1.1 we deduce that

$$\frac{d}{dt}(y(t+T) - y(t)) = -(y(t+T) - y(t)) \tag{1.22}$$

and therefore, using $g \equiv -1$ in (1.8), we have that the exists $k \in \mathbb{R}$ such that

$$y(t+T) - y(t) = k e^{-t} \xrightarrow[t \to \infty]{} 0. \tag{1.23}$$

□

Thus, the solution tends to a periodic regime. Note that the right side of the equality is positive, meaning that $y(t+T) > y(t)$ for all $t \geq 0$, implying that $y(t)$ oscillates increasingly until the periodic regime.

**Remark 1.2.** In particular, for a periodic pulse as in (1.1) and initial datum $y(0) = 0$, we can compute $y(T)$ from equation (1.21)

$$y(T) = \int_0^T F(\gamma(r)) e^{-T+r} dr = e^{-T} F(A_0 T)\left(e^{d_0} - 1\right) \tag{1.24}$$

and therefore from (1.9), we have that

$$y(t+T) - y(t) = y(T) e^{-t} = e^{-t-T} F(A_0 T)\left(e^{d_0} - 1\right). \tag{1.25}$$

1.2.2. **Time scales.** We now analyze the time $\tau$ (the time to reach steady state when stimulated with a constant input conserving dose) , the time $\tau_p$ (the time it takes the system to *reach the periodic solution*) and compare them. When the system is forced with a constant input $\gamma(t) \equiv A_0 d_0$, the solution of equation (1.21) becomes

$$y(t) = F\left(A_0 d_0\right)\left(1 - e^{-t}\right)$$

and therefore the system takes time $\tau = 1$ to reach $(1 - e^{-1})$ times the steady state $F(A_0 d_0)$.

In order to compute $\tau_p$ we assume that the system is stimulated with a periodic square input $\gamma$ as defined in (1.1). Given $\varepsilon > 0$ small we search for a time $\tau_p$ for which $|y(t+T) - y(t)| < \varepsilon$ for all $t > \tau_p$.

Note that, from (1.25), $y(t+T) - y(t) = y(T)e^{-t} \leq y(T)$, if we take $\varepsilon > 0$ such that $y(T) < \varepsilon$ we would have that $y(t+T) - y(t) < \varepsilon$ for all $t > 0$.

We take $\varepsilon > 0$ small enough such that the solutions $y(t)$ and $y(t+T)$ at $t = 0$ are distant with respect to $\varepsilon$, that is $y(0+T) - y(0) = y(T) > \varepsilon$. From (1.25), this last condition is equivalent to

$$e^{-T} F\left(A_0 T\right)\left(e^{d_0} - 1\right) > \varepsilon. \tag{1.26}$$

Then, the time $\tau_p$ that the solution takes to sastisfy $|y(t+T) - y(t)| < \varepsilon$ for all $t > \tau_p$ is given by

$$\tau_p(T) = \ln\left(F\left(A_0 T\right)\left(e^{d_0} - 1\right)e^{-T}\varepsilon^{-1}\right). \tag{1.27}$$

Note that for each $T$ such that condition (1.26) is satisfied, we have that $\tau_p > 0$.

If $F$ is a function such that $F(A_0 T)e^{-T} \leq M$ for all $T > 0$ and $(e^{d_0} - 1)M\varepsilon^{-1} < e$ then

$$\tau_p(T) < \tau = 1 \quad \text{for all} \quad T.$$

Moreover, if $\lim_{T\to\infty} F(A_0 T)e^{-T} = 0$ then for all $\varepsilon > 0$ $\tau_p(T) \to -\infty$.

1.2.3. $\tau_{p,max}$ **for** $T$ **bounded.** Conversely, if $T$ **is bounded**, for instance assume $d_0 < T < T_{max}$ and since$F$ is an increasing function we have the following uniform in $T$ inequalities

$$F\left(A_0 d_0\right) e^{-T_{max}} < F\left(A_0 T\right) e^{-T} < F(A_0 T_{max})e^{-d_0}. \tag{1.28}$$

From the left hand side inequality, if we assume that the constants $A_0, d_0, T_{max}$ satisfy

$$\varepsilon < F\left(A_0 d_0\right)\left(e^{d_0} - 1\right)e^{-T_{max}} \tag{1.29}$$

we would have that $\tau_p(T) > 0$ for all $T$. And if we assume that

$$\varepsilon < F\left(A_0 d_0\right)\left(e^{d_0} - 1\right)e^{-T_{max}}e^{-1}. \tag{1.30}$$

we would deduce from (1.27) that $\tau_p(T) > 1 = \tau$ for all $T$. Moreover, from the right hand side inequality (1.28) we derive

$$\tau_p(T) < \ln\left(F(A_0 T_{max})(1 - e^{-d_0})\varepsilon^{-1}\right) := \tau_{p,max} \tag{1.31}$$

independent of $T \in [d_0, T_{max}]$. Putting all together we have that if $T \in [d_0, T_{max}]$ and condition (1.30) is satisfied then

$$1 = \tau < \tau_p(T) < \tau_{p,max} \text{ for all } T \in [d_0, T_{max}].$$

1.3. **Coupled signaling components without feedback connections.**

1.3.1. **The solution is asymptotically periodic.** Finally we prove that the solution of model (1.4) is an asymptotically periodic function.

**Proposition 1.3.** *Given $F_1, F_2 : \mathbb{R} \to \mathbb{R}$ $C^1$ functions $\gamma(t)$ a $T$-periodic function and $F_2$ is increasing. Then the solution of equation* (1.4) *is asymptotically a $T$-periodic function.*

*Proof.* From proposition 1.2 there exists a constant $c_1$ such that

$$y_1(t+T) - y_1(t) = c_1 e^{-t} \xrightarrow[t\to\infty]{} 0$$

Also, since

$$\tau_{21}\frac{d}{dt}(y_2(t+T) - y_2(t)) = F_2(y_1(t+T)) - F_2(y_1(t)) - (y_2(t+T) - y_2(t)).$$

there exists and a constant $c_2$ such that

$$\begin{aligned} y_2(t+T) - y_2(t) &= e^{-t/\tau_{21}}\left(c_2 + \int_0^t \left(F_2(y_1(s+T)) - F_2(y_1(s))\right) e^{s/\tau_{21}} ds\right) \\ &= e^{-t/\tau_{21}}\left(c_2 + \int_0^t \left(F_2(y_1(s) + c_1 e^{-s}) - F_2(y_1(s))\right) e^{s/\tau_{21}} ds\right) \\ &= e^{-t/\tau_{21}}\left(c_2 + \int_0^t F_2'(\xi(s)) c_1 e^{-s} e^{s/\tau_{21}} ds\right) \end{aligned}$$

where $\xi(s)$ is an intermediate point between $y_1(s)$ and $y_1(s) + c_1 e^{-s}$. Since $F_2$ is increasing and $\xi(s)$ is bounded we have that $0 < F_2'(\xi(s)) \leq M$ for all $s \geq 0$. Then

$$\int_0^t F_2'(\xi(s)) e^{-s} e^{s/\tau_{21}} ds \leq M \int_0^t e^{s(1/\tau_{21}-1)} ds.$$

For $\tau_{21} = 1$, we obtain that

$$\int_0^t e^{s(1/\tau_{21}-1)} ds = t,$$

and for $\tau_{21} > 0$, $\tau_{21} \neq 1$

$$\int_0^t e^{s(1/\tau_{21}-1)} ds = \frac{e^{t(1/\tau_{21}-1)} - 1}{1/\tau_{21} - 1}$$

Thus, in either case we have proved that

$$|y_2(t+T) - y_2(t)| \leq e^{-t/\tau_{21}}(c_2 + M\int_0^t e^{s(1/\tau_{21}-1)} ds) \xrightarrow[t\to\infty]{} 0.$$

□

### 2. The signaling component with a simplified linear description response to frequency depends on the concavity of the function $F$

In this section we show how the frequency response of the observable $G_S$ (sliding window) depends on the concavity of the function $F$. We consider three types of monotonic increasing functions: concave, convex and sigmoidal ones. A real-valued function defined in an interval is called convex/concave if the line segment between any two points on the graph of the function lies above/below the graph. For a derivable function of a single variable, a function is convex/concave on an interval if and only if the tangent line on every point of the interval lies below/above the graph. Moreover, for a twice differentiable function of a single variable, if the second derivative is always greater/lower than zero for its entire domain then the function is convex/concave.

For a concave function satisfying $F(0) = 0$ it holds: $F(\lambda x) \geq \lambda F(x)$ for all $\lambda \in [0,1]$. Such functions are also called sublinear. Analogously, for a convex function satisfying $F(0) = 0$ it holds: $F(\lambda x) \leq \lambda F(x)$ for all $\lambda \in [0,1]$ and such functions are also called supralinear. A linear function such that $F(0) = 0$ is the one that satisfies the equality $F(\lambda x) = \lambda F(x)$ for all $\lambda \in [0,1]$. A sigmoidal function $F(x)$ is concave for $x < x_0$ and convex for $x > x_0$ for some $x_0$.

We consider the following lemma that is useful for the analytical study of the frequency response of this system.

**Lemma 2.1.** *Given $F : [0,c] \to \mathbb{R}$ a continuous function, derivable on $(0,c)$ such that $F(0) = 0$ and $F(x) > 0$ for all $x > 0$.*

*(1) If $F$ is linear in $[0,c]$, then $F'(x)x = F(x)$ for all $x \in [0,c]$.*

*(2) If $F$ is concave in $[0,c]$ then $\dfrac{F'(x)x}{F(x)} \leq 1$ for all $x \in (0,c)$. Moreover, the inequality is strict for $F$ strictly concave.*

*(3) If $F$ is convex in $[0,c]$ then $\dfrac{F'(x)x}{F(x)} \geq 1$ for all $x \in (0,c)$. Moreover, the inequality is strict for $F$ strictly convex.*

*Proof.* The proof is straightforward for a linear function satisfying $F(0) = 0$. Since $F$ is a derivable function on $[0,c]$ and $F(0) = 0$, the tangent line on each $x \in [0,c]$ is given by $L(y) = F(x) + F'(x)(y-x)$. For concave functions, we have that $F(y) \leq L(y)\ \forall y \in [0,c]$, in particular for $y = 0$ obtaining that $0 \leq F(x) - xF'(x)$. Using the positivity of $F$ we get the desired result. Analogously we obtain the corresponding result for convex functions. □

Using Lemma 2.1 it is possible to demonstrate which is the frequency response for the observable $G_S$ depending on the function's concavity, where:

$$G_S(T) = \frac{1}{T}\int_S^{S+T} y(t)dt.$$

Using the definition of $\gamma$ given in (1.1) and considering that the dose is conserved by amplitude compensation, from the solution of equation (1.21) given in (1.21), we compute the average

$$G_S(T) = \frac{1}{T}\left(\int_S^{S+T}\int_0^S F(\gamma(r))e^{-t+r}drdt + \int_S^{S+T}\int_S^t F(\gamma(r))e^{-t+r}drdt\right).$$

For the first term we have

$$\int_S^{S+T}\int_0^S F(\gamma(r))e^{-t+r}drdt = e^{-S}(1-e^{-T})\left(\int_0^S F(\gamma(r))e^r dr\right).$$

For the second term, changing the order of the variables (Fubini's Theorem), we obtain

$$\begin{aligned}\int_S^{S+T}\left(\int_S^t F(\gamma(r))e^{-t+r}dr\right)dt &= \int_S^{S+T} F\left(\gamma(r)\right)\left(1-e^{r-S-T}\right)dr\\ &= F\left(A_0T\right)d_0 - e^{-S-T}\left(\int_0^{S+T} F\left(\gamma(r)\right)e^r dr - \int_0^S F\left(\gamma(r)\right)e^r dr\right).\end{aligned}$$

Combining both terms we deduce

$$G_S(T) = \frac{1}{T}\left(F\left(A_0T\right)d_0 + e^{-S}\int_0^S F(\gamma(r))e^r dr - e^{-S-T}\int_0^{S+T} F(\gamma(r))e^r dr\right).$$

Using that $\gamma$ is $T-$ periodic

$$\begin{aligned}\int_0^{S+T} F(\gamma(r))e^r dr &= \int_0^T F(\gamma(r))e^r dr + \int_T^{S+T} F(\gamma(r))e^r dr\\ &= F\left(A_0T\right)\left(e^{d_0}-1\right) + e^T\int_0^S F(\gamma(r))e^r dr,\end{aligned}$$

which leads to the following analytical expression for the observable $G_S$ as a function of the input's period $T$:

$$G_S(T) = \frac{1}{T}F\left(A_0T\right)\left(d_0 - e^{-T-S}(e^{d_0}-1)\right). \tag{2.1}$$

In order to evaluate for which periods (or equivalently frequencies) the observable is optimized, we should study the derivative with respect to $T$:

$$\begin{aligned}G'_S(T) = &\frac{1}{T^2}F'\left(A_0T\right)A_0T\left[d_0 - e^{-(T+S)}(e^{d_0}-1)\right]\\ &-\frac{1}{T^2}F\left(A_0T\right)\left[d_0 - e^{-(T+S)}(e^{d_0}-1)(T+1)\right].\end{aligned} \tag{2.2}$$

**Remark 2.1.** For $S$ big enough, where the system is in the periodic regime, from (2.1) the average $G_S(T) \sim \frac{d_0}{T} F(A_0 T)$, which is the same expression for the average used by Fletcher [?] when $F(0) = 0$. Moreover, from (2.2), the derivative

$$G'_S(T) \sim \frac{F(A_0 T)}{T^2} d_0 \left( \frac{F'(A_0 T) A_0 T}{F(A_0 T)} - 1 \right). \tag{2.3}$$

Using this approximation for the derivative $G'_S(T)$, we obtain that the sign of this expression is established by the concavity of the production function $F$. As the frequency is the inverse of the period T, and we study the sign of the derivative in terms of the period, when this derivate is positive (negative) we will have a lowpass (highpass) filter. More precisely, when $F$ is a concave function from item (2) of lemma 2.1 we have that $G'_S(T) < 0$ and therefore the response vs. frequency curve is a highpass filter. On the other hand, when $F$ is a convex function from item (1) of lemma 2.1 we have that $G'_S(T) > 0$ and therefore the response vs. frequency curve is a lowpass filter. Finally when $F$ is linear, $G'_S(T) \sim 0$.

The following lemma will be usefull for the characterization of the sign of $G'_S(T)$.

**Lemma 2.2.** *Given $S > 0$, we define the function $C_S : [d_0, +\infty) \to \mathbb{R}$ as follows*

$$C_S(T) := \frac{d_0 - e^{-(T+S)}(e^{d_0} - 1)(T+1)}{d_0 - e^{-(T+S)}(e^{d_0} - 1)} = 1 - \frac{e^{-(T+S)}(e^{d_0} - 1)T}{d_0 - e^{-(T+S)}(e^{d_0} - 1)}. \tag{2.4}$$

*We have that:*

*(1) $C_S(T) < 1$ for all $T \geq d_0$.*

*(2) $\lim_{T\to+\infty} C_S(T) = 1$ for all $S > 0$ and $\lim_{S\to+\infty} C_S(T) = 1$ for all $T > d_0$.*

*(3) $C'_S(T) > 0 \iff (1-T)e^{(T+S)} < \frac{(e^{d_0}-1)}{d_0}$.*

*(4) $G'_S(T) < 0 \iff \frac{F'(A_0T)A_0T}{F(A_0T)} < C_S(T)$.*

*(5) $G'_S(T) > 0 \iff \frac{F'(A_0T)A_0T}{F(A_0T)} > C_S(T)$.*

*Proof.* The proofs of items (1)-(2) are straightforward computations. Item (3) is a direct consequence of the derivative given by

$$C'_S(T) = -\frac{e^{-(T+S)}(e^{d_0} - 1)}{\left[d_0 - e^{-(T+S)}(e^{d_0} - 1)\right]^2} \left[(1-T)d_0 - e^{-(T+S)}(e^{d_0} - 1)\right].$$

For the last two items we used that for $0 \leq d_0 < T$ and $S > 0$, we have $d_0 - e^{-(T+S)}\left(e^{d_0} - 1\right) > d_0 - 1 + e^{-d_0} > 0$. □

**Lemma 2.3.** *For $F$ a convex or linear function satisfying the conditions of lemma 2.1, the derivative $G'_S$ is always positive, concluding that the response vs. frequency curve is a lowpass filter for any $S \geq 0$.*

*Proof.* Combining item (3) from Lemma 2.1 with item (1) from Lemma 2.2, from item (5) in Lemma 2.2 we obtain the desired result. In the case of linear functions, using item (1) from Lemma 2.1, we arrive to the same conclusion. □

For concave functions, we can find different responses in the transient phase. For instance, we will consider two different families of concave functions given in the following examples.

**Example 2.1.** Consider the family of concave functions $F_\alpha(x) = x^\alpha$, with $0 < \alpha < 1$ which satisfies conditions of lemma 2.1. In this case for $x > 0$

$$\frac{F'_\alpha(x)x}{F_\alpha(x)} = \frac{\alpha x^{\alpha-1}x}{x^\alpha} = \alpha, \tag{2.5}$$

which is independent of the paremeters $d_0$, $S$ and $A_0$. If we assume $S = 0.16, d_0 = 0.1$ and $A_0$ arbitrary, from Fig. 1, we see that:

- For $\alpha = 0.7$, there exists $T_1$ such that the horizontal line with constant value 0.7 (in blue) is bigger than $C_S(T)$ for all $T \in [d_0, T_1)$ and smaller than $C_S(T)$ for all $T > T_1$. Then, from items (4) and (5) in Lemma 2.2 and equation (2.5) we have that $G'_S(T) > 0$ for all $t \in [d_0, T_1)$ and $G'_S(T) < 0$ for all $t > T_1$. Thus, we have a frequency preference.
- For $\alpha = 0.45$ with a similar analysis we obtain that there are two changes in the sign of $G'_S(T)$ since the horizontal line with constant value 0.45 (pink line) crosses the graph of $C_S$ twice, obtaining an antiresonance and resonance behavior.
- For $\alpha = 0.2$, similarly we obtain a highpass filter since the horizontal line with constant value 0.2 (in red) is always smaller that $C_S(T)$.

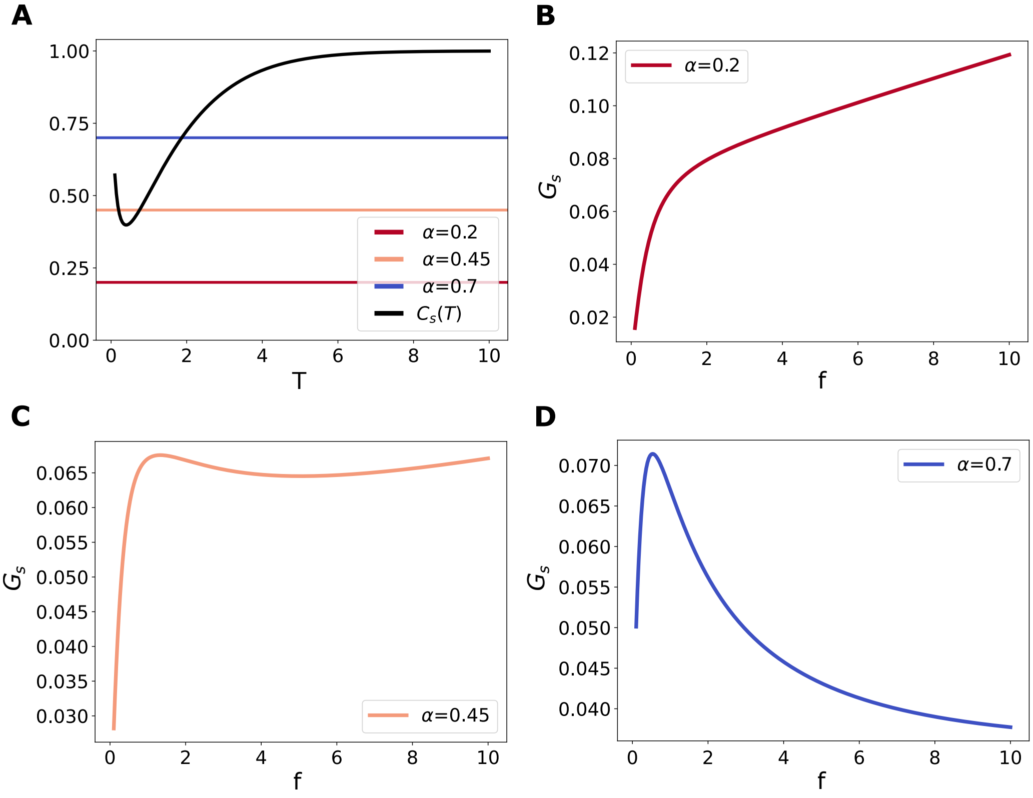


FIGURE 1. **Frequency response for the signaling component model with the concave production function $F(x) = x^{\alpha}$. A.** $C_S(T)$ in black. The horizontal lines represent the values of $\alpha = 0.2, 0.45, 0.7$. **B-D.** $G_S$ as a function of frequency as the parameter $\alpha$ takes different values with $A_0 = 1, d_0 = 0.1$ and $S = 0.16$. **B**: $\alpha = 0.2$, **C:** $\alpha = 0.45$, **D**: $\alpha = 0.7$.

**Example 2.2.** Consider the family of concave functions $F_k(x) = \frac{x}{x+k}$, with $k > 0$. A straightforward computation gives

$$\frac{F'_k(x)x}{F_k(x)} = \frac{k}{x+k}. \tag{2.6}$$

We define the function $R_S(T) = \dfrac{A_0 T C_S(T)}{1 - C_S(T)}$, where $C_S(T)$ is the function defined in (2.4). From items (4) and (5) of lemma 2.2 and equation (2.6) we deduce that

$$G'_S(T) > 0 \iff \frac{k}{A_0 T + k} > C_S(T) \text{ and } G'_S(T) < 0 \iff \frac{k}{A_0 T + k} < C_S(T)$$

and thus

$$G'_S(T) > 0 \iff k > R_S(T) \text{ and } G'_S(T) < 0 \iff k < R_S(T).$$

Note that since $\lim_{S \to +\infty} R_S(T) = +\infty$ for all $T$, when $S$ is big enough, we obtain that $k < R_S(T)$ for all $T$ and therefore we have a highpass filter as we have shown in Remark 2.1.

Moreover, since

$$R_S'(T) = \frac{A_0}{(e^{d_0}-1)}(1 - e^{d_0} + d_0 e^{S+T}) > 0 \text{ for all } T > d_0 \text{ and } S > 0 \tag{2.7}$$

we have that $R_S(T)$ is a strictly increasing function. Considering the parameters $A_0, d_0$ and $S$ the same as in the latter example, we deduce (see Fig. 2):

- For $k < R_S(d_0) \sim 0.13$ (red line), we have $k < R_S(T)$ for all $T \in [d_0, \infty]$. Therefore we have a highpass filter.
- For $k > R_S(d_0)$, since $\lim_{T\to+\infty} R_S(T) = +\infty$ for all $S > 0$, we can take $T_1 = R_S^{-1}(k)$ for which $k > R_S(T)$ for all $T \in [d_0, T_1)$ and $k < R_S(T)$ for all $T > T_1$ (blue and pink lines). Thus, we have frequency preference.

In Fig. 2 we show the average $G_S(T)$ for the different values of $k$. Note that for $k > R_S(d_0)$, the value where the maximum of $G_S$ (as a function of the frequency) is attained is deacreasing with respect to $k$.

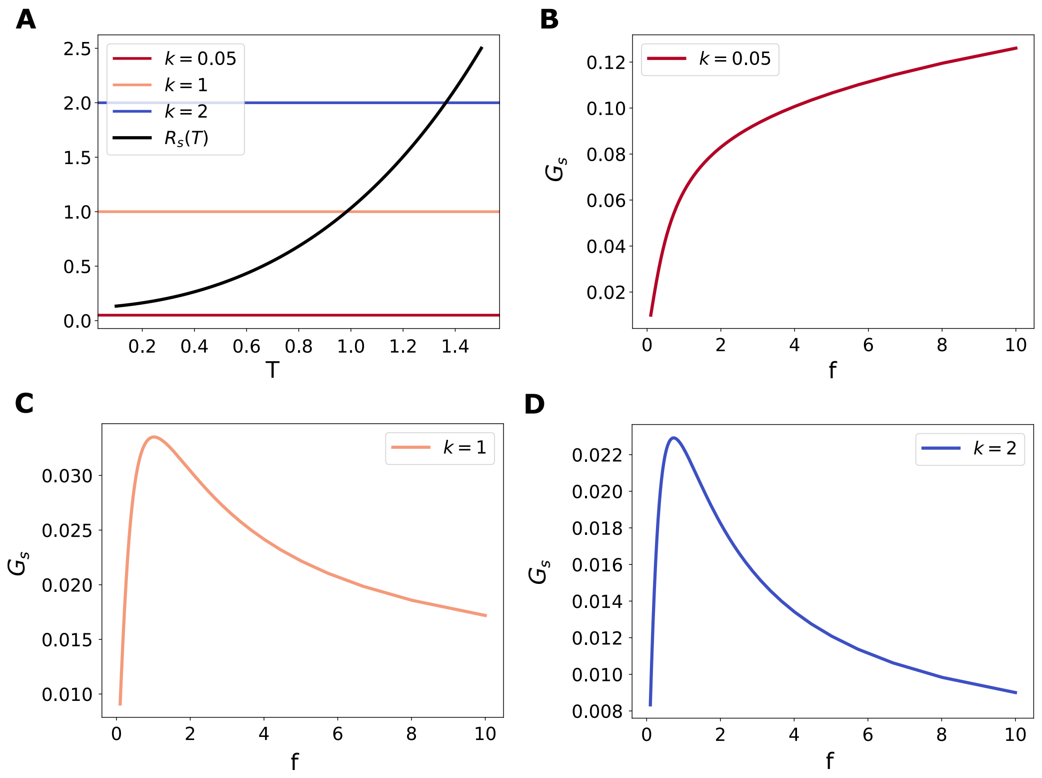


FIGURE 2. **Frequency response for the signaling component model with the concave production function** $F(x) = x/(x+k)$**. A.** $R_S(T)$ in black. The horizontal lines represent the values of $k = 0.05, 1, 2$. **B-D.** $G_S$ as a function of frequency as the parameter $k$ takes different values with $A_0 = 1, d_0 = 0.1$ and $S = 0.16$. **B**: $k = 0.05$, **C:** $k = 1$, **D**: $k = 2$.

We now assume that a production function $F$ is sigmoidal, being convex in $[0, x^*]$ and concave in $[x^*, \infty)$ for some $x^* > 0$. From 2.3, for the values of $T \geq d_0$ such that $A_0 T < x^*$ we will have that the average $G_S(T)$ is increasing. After that we cannot expect that the average behaves like for concave functions since the production function is not zero on $x^*$..

Next, we will show a family of sigmoidal functions for which we observe different behaviors.

**Example 2.3.** We consider the family of functions $F_k(x) = \dfrac{x^2}{x^2+k}$. These functions change its concavity at $x^* = \sqrt{k/3}$ from convex to concave. We have the following equality for $x > 0$

$$\frac{F_k'(x)x}{F_k(x)} = \frac{2k}{x^2+k}. \tag{2.8}$$

We define the function $Q_S(T) = \dfrac{(A_0T)^2\, C_S(T)}{2 - C_S(T)}$, where $C_S(T)$ is the function defined in (2.4). From items (4) and (5) of lemma 2.2 and equation (2.8) we deduce that

$$G'_S(T) > 0 \Longleftrightarrow \frac{2k}{(A_0T)^2 + k} > C_S(T) \text{ and } G'_S(T) < 0 \Longleftrightarrow \frac{2k}{(A_0T)^2 + k} < C_S(T)$$

and thus

$$G'_S(T) > 0 \Longleftrightarrow k > Q_S(T) \text{ and } G'_S(T) < 0 \Longleftrightarrow k < Q_S(T).$$

We compute the derivative of the function $Q_S$ and obtain

$$Q'_S(T) = \frac{2A_0^2T}{(2 - C'_S(T))^2}\frac{(e^{d_0} - 1)(1 + T - T^2)e^{-T-S} - d_0}{e^{-T-S}(e^{d_0} - 1) - d_0}.$$

Since $e^{-T-S}(e^{d_0}-1)-d_0 < 0$, we have that $Q'_S(T) > 0$ if and only if $(e^{d_0}-1)(1+T-T^2)e^{-T-S}-d_0 < 0$. Thus,

$$Q'_S(T) > 0 \Leftrightarrow (1 + T - T^2)e^{-T-S} < \frac{d_0}{e^{d_0} - 1}.$$

Note that the function $h(T) = (1 + T - T^2)e^{-T}$ decreases in $[0, 3]$, increases in $[3, +\infty)$, $h(T) > 0$ for $T \in [0, \frac{1}{2} + \sqrt{5}]$ and $h(T) < 0$ for $T > \frac{1}{2} + \sqrt{5}$. Therefore, for $d_0 < \frac{1}{2} + \sqrt{5}$, $h(T) < h(d_0)$ for all $T > d_0$. Then, we deduce that if we choose $d_0 < \frac{1}{2} + \sqrt{5}$ and $S$ such that

$$(1 + d_0 - d_0^2)e^{-d_0} < e^S \frac{d_0}{e^{d_0} - 1} \tag{2.9}$$

we will have that $Q_S$ is increasing. Then, for parameters $d_0$ with $0 < d_0 < \frac{1}{2} + \sqrt{5}$ and $S$ such that condition (2.9) is satisfied, $Q_S(T)$ is a strictly increasing function. Also $\lim_{T\to+\infty} Q_S(T) = +\infty$ for all $S > 0$. Therefore we can choose different values of $k$ to obtain (see Fig. 3):

(1) a highpass filter, if $k < Q_S(d_0) \sim 0.000999$, for instance the red line.
(2) resonance, if $k > Q_S(d_0)$, for instance the blue or pink lines.

In Fig. 3 we show the average $G_S(T)$ for the different values of $k$. Note that for $k > Q_S(d_0)$, the value where the maximum of $G_S$ as a function of frequency is attained is decreasing with respect to $k$.

### 3. Coupled covalent modification cycle does not generate feedback over the ligand-receptor system

In this section, we demonstrate that a covalent modification cycle coupled to a ligand-receptor system as considered in the main text (see Methods Eq. M6-M18) does not generate feedback over the first level. The stimulus is the free ligand amount $L$. The observable over the first node is calculated over $y = RL + RLS$. The time derivative is:

$$\frac{dy}{dt} = \frac{dRL}{dt} + \frac{dRLS}{dt} \tag{3.1}$$

Usign the differential equations M11 and M13 from Methods we obtain:

$$\frac{dy}{dt} = k_+R.L - k_-.RL - a_1.RL.S + (d_1 + k_1).RLS + k_+.RS.L - k_-.RLS - (d_1 + k_1).RLS \tag{3.2}$$

Then, using the conservation law for the receptor $R_T = R + RL + RS + RLS$ and the definition of $y$, we obtain:

$$\frac{dy}{dt} = k_+L(R_T - y) - k_-y \tag{3.3}$$

The resulting equation is identical to the one for the isolated ligand-receptor system (Eq.R1), which implies that there is no feedback from the second node (CMC) to the first one. Thus, the activity of the first node and its response as a function of the input frequency is described in the first section of Results.

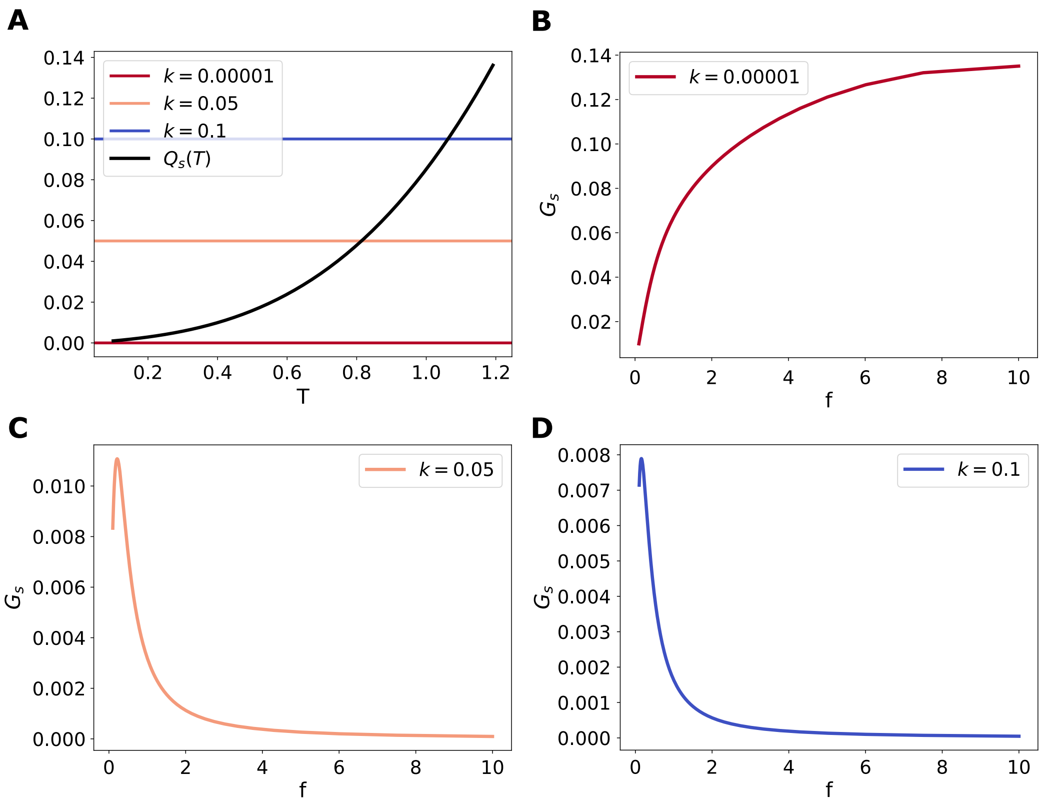


FIGURE 3. **Frequency response for the signaling component model with the sigmoidal production function** $F(x) = x^2/(x^2+k)$**. A.** $Q_S(T)$ in black. The horizontal lines represent the values of $k = 0.00001, 0.05, 0.1$. **B-D.** $G_S$ as a function of frequency as the parameter $k$ takes different values with $A_0 = 0.5, d_0 = 0.1$ and $S = 0.16$. **B**: $k = 0.00001$, **C:** $k = 0.05$, **D**: $k = 0.1$.

### 4. Additional results for the ligand-receptor model

4.1. **Transient frequency preference is also observed for a different periodic stimulation.** The ligand-receptor system exhibited a transient frequency preference when stimulated with a train of square pulses. In order to elucidate if this phenomenon is robust, we explored a different stimulation, using a sinusoidal function. The two observables described in Methods ($G_S$ and $H_S$) were computed for a range of frequencies. Results are shown in Fig. 4. Frequency preference is observed for the temporal average $G_S$ for lower values of $S$.

4.2. **Dose dependence in frequency response in the ligand-receptor system.**

4.2.1. *Dose dependence in the pseudo-periodic regime.* We analyzed the influence of the input parameters (amplitude, duration, and dose) in the ligand receptor frequency response. In the main text, this analysis was performed for a time window such that the response is in the transient regime ($S = 0.1$) and using the conservation scheme by amplitude compensation. Here we study how the amplitude, the duration and the mean dose of the input (a train of square pulses) shape the frequency response in the observable $G_S$ in the periodic regime. As well as in the main text, we consider $T_0 = 1$. Results are shown in Fig. 5.

In Fig. 5A the duration of the pulses is fixed, $d_0 = 0.1$, while $A_0$ takes different values, resulting in different mean input doses $d_0A_0$. In all the cases, the frequency response in the periodic regime is a high pass filter. As the amplitude is increased, the observable $G_S$ reaches a higher value. A similar result is found by fixing the reference amplitude $A_0$ and increasing the pulse duration $d_0$, thus the mean dose of the stimuli is increased correspondingly (Fig. 5B). In Fig. 5C, both $d_0$ and $A_0$ are varied while keeping the mean input $d_0A_0$ constant. Only in this case, the frequency response is almost identical for all the scenarios studied, suggesting that the mean input dose is

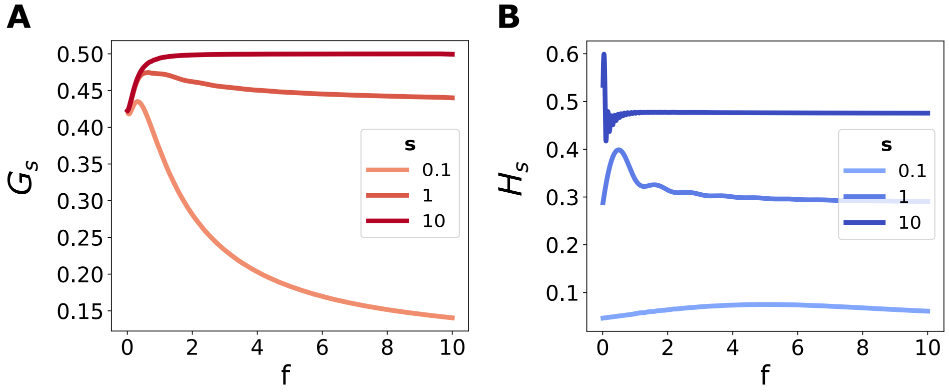


FIGURE 4. **Transient frequency preference response for the ligand-receptor system with a sinusoidal stimulation.** Observables $G_S$ (A) and $H_S$ (B) as a function of frequency, for different time windows, indicated by the parameter $S$. Lighter green indicates low values of $S$, recalling a transient phase. Parameters: $A_0 = 10$.

key in the frequency response of the system, as well as the temporal window where it is computed.

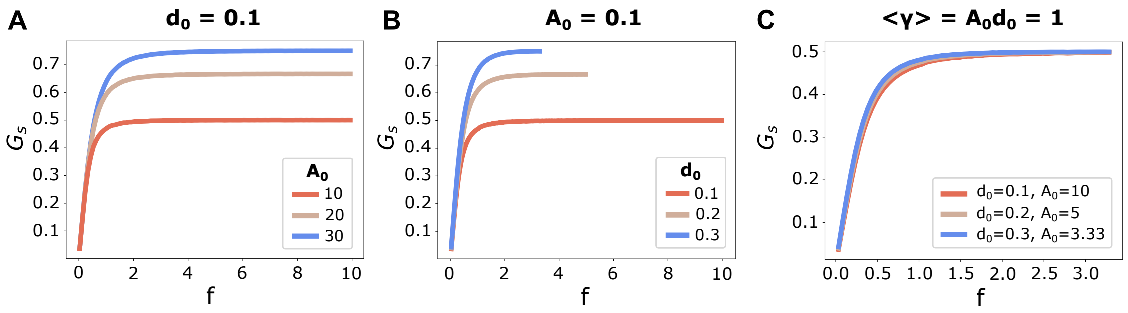


FIGURE 5. **Input parameters influence on the frequency response for the ligand-receptor system.** Observable $G_S$ as a function of frequency, measured in the periodic regime, with time $S = 10$. Dose was conserved by varying the pulses amplitude. **A.** Fixed reference duration $d_0 = 0.1$ and varying the reference amplitude $A_0$, thus the mean dose is $\langle\gamma\rangle = A_0 d_0$. **B.** Fixed amplitude $A_0 = 10$, for each curve a different dose given by $d_0$. **C.** Fixed dose for all the curves, varying $d_0$ and $A_0$. Each curve was simulated until the maximum frequency, given by $f_{cut} = 1/d_0$.

4.2.2. *Dose dependence in the transient regime with the dose conservation by duration compensation.* In this section we study the influence of the stimulus parameters in the transient frequency response when dose conservation is achieved by modifying the duration of the pulses. Results are shown in Fig. 6. As $A_0$ increases, the preferred frequency $f_{G,max}$ shifts towards higher values and $G_{max}$ also increases. A similar result is found by fixing the reference amplitude $A_0$ and increasing the pulse duration $d_0$. In Fig. 6, both $d_0$ and $A_0$ are varied while keeping the mean input dose constant. Only in this case, $f_{G,max}$ and $G_{max}$ remain practically constant, suggesting that the preferred frequency and maximum value for the sliding window are controlled by the mean input dose. In Fig.6 D and E we explore the influence of the mean input dose on the frequency preference response. The preferred frequency $f_{G,max}$ and the maximum value $G_{max}$ are plotted as functions of the mean dose. The preferred frequency $f_{G,max}$ varies with the mean input dose in a similar way for the different values of $d_0$. Together, these studies support the underlying hypothesis that the fundamental parameter to understand the transient frequency response in a ligand-receptor system is the input mean dose.

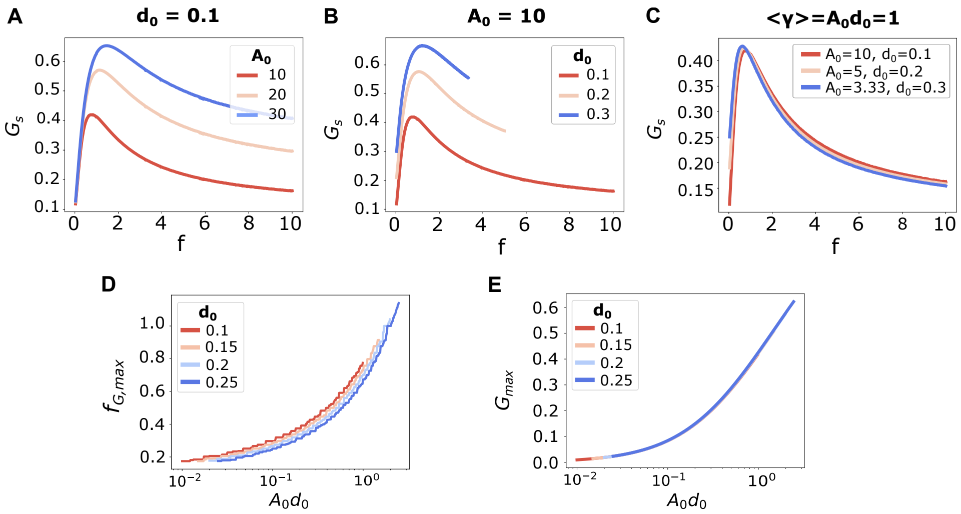


FIGURE 6. **Input parameters influence on the frequency response.** Observable $G_S$ as a function of frequency, measured at a transient time $S = 0.1$. Dose was conserved by varying the pulses duration. **A.** Fixed reference duration $d_0 = 0.1$, period $T_0 = 1$ and varying the amplitude $A_0$, thus the mean dose is $\langle\gamma\rangle = A_0 d_0$. **B.** Fixed amplitude $A_0 = 10$, for each curve a different dose given by $d_0$. **C.** Fixed dose for all the curves, varying $d_0$ and $A_0$. **D.** Preferred frequency for different frequency curves of the observable $G_S$. $f_{G,max}$ as a function of the mean dose $A_0 d_0$. **E.** Maximum observable $G_{max}$ as a function of the mean dose $A_0 d_0$. The duration $d_0$ was fixed as the amplitude $A_0$ varies. Different durations $d_0$ are indicated in colors.

### 5. Additional results for signaling components with a simplified linear description

5.1. **Expanding window results.** In this section, we study the frequency response for the signaling component with a simplified linear description for the expanding window. We examine the observable $H_S$ as a function of the input frequency, considering an amplitude compensation scheme to maintain dose conservation.

In Fig. 7 we show the results for the observable for functions $F$ with representative shapes and for a different values of $S$. For this observable, there is a minimum frequency given by $f_{min} = 1/S$ to have at least one oscillation in the time window. Thus, for shorter initial times the range of frequencies is narrower.

As in the results shown in the main text, the frequency response depends both of the nonlinearity of the function and the time window where it is computed. Local oscillations are observed due to small variations in the mean dose over the time interval $[0, S]$ where the observable is computed, since it may not contain a complete number of periods.

### 6. Additional results for the MAPK cascade model

In this section we study the influence of the input parameters in the transient frequency response. For all the results in this section, the parameter set was taken from [2] and the observable $G_S$ as a function of frequency is computed from an initial time $S = 0.1min$.

In Fig. 8 we analyze how the observable $G_S$ in each level of the cascade is shaped as we change the duration of the input reference pulse, for both dose conservation protocols. For the three levels, as the duration of the pulse is incresead the prefered frequency shiftes to lower values. Moreover, the preference is more pronounced.

In Fig. 9 we study how the amplitude of the reference input impacts on the frequency response in each level of the cascade for both dose conservation schemes. In both cases, the prefered frequency and its corresponding value $G_{max}$ is higher as the reference amplitude is increased for all the three levels.

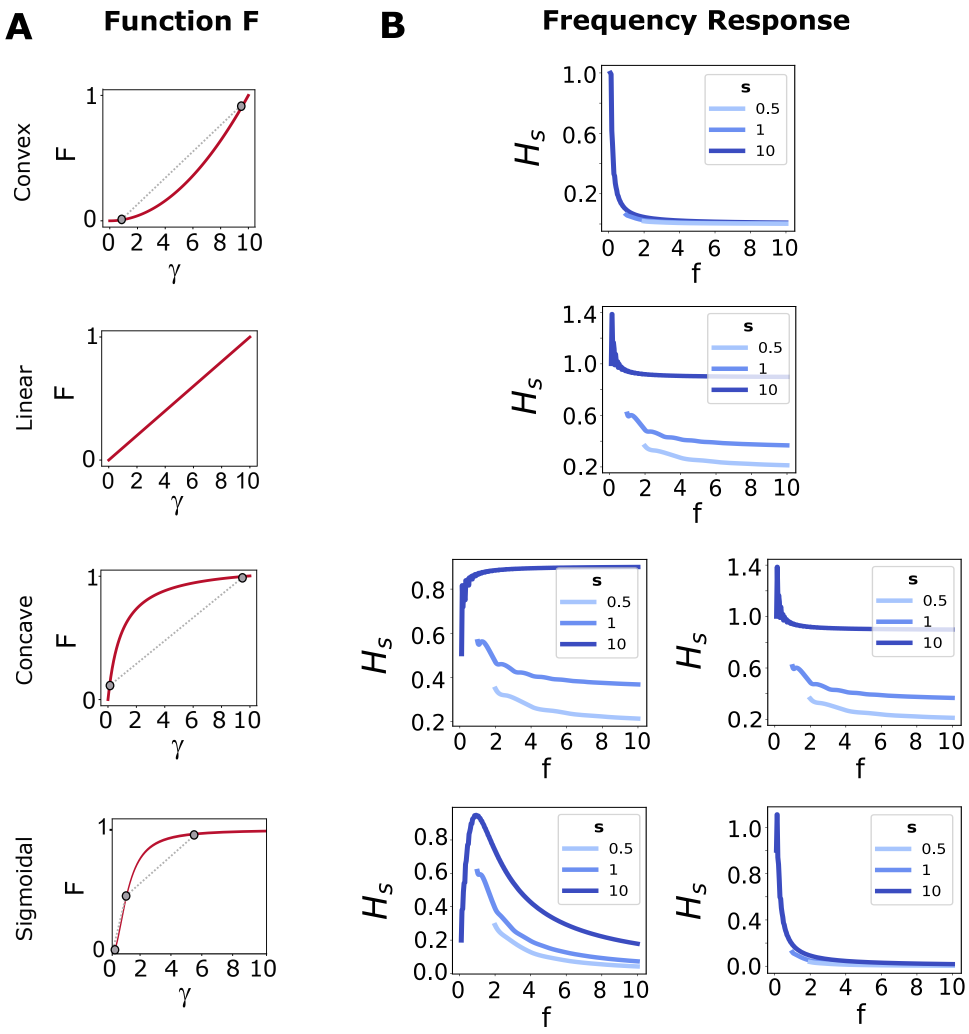


FIGURE 7. **The frequency response for a one-dimensional signalling component depends on the concavity of the function** $F(\gamma)$. The system is stimulated with a periodic train of square pulses, using an amplitude compensation protocol to conserve input dose. Input parameters: $A_0 = 1, d_0 = 1, T_0 = 1$. The observable computed is the expanding window $H_S$, as described in Methods, normalized by the maximum value for the sliding window results $G_S$ (Fig. 4 in the Main Text). **A.** Examples for the four possible concavities for function $F(\gamma)$. **B.** Frequency response for the four concavities (rows) for different values of $S$, earlier times in lighter lines. For concave and sigmoidal functions, different responses are achieved for different parameter values. The functions selected for this figure are: convex $F(\gamma) = \gamma^2$, linear $F(\gamma) = \gamma$, concaves $F(\gamma) = \gamma/(10 + \gamma)$ (left) and $F(\gamma) = \gamma/(1000 + \gamma)$ (right) and sigmoidals $F(\gamma) = \gamma^2/(1 + \gamma^2)$ (left) and $F(\gamma) = \gamma^2/(100 + \gamma^2)$ (right).

In Fig. 10 we compute the frequency response for different combinations of amplitude and duration but with the same dose. We can see that the prefered frequency value takes lower values and the maximum observable value is higher as the duration is increased and the amplitude drecreases, even the dose is fixed.

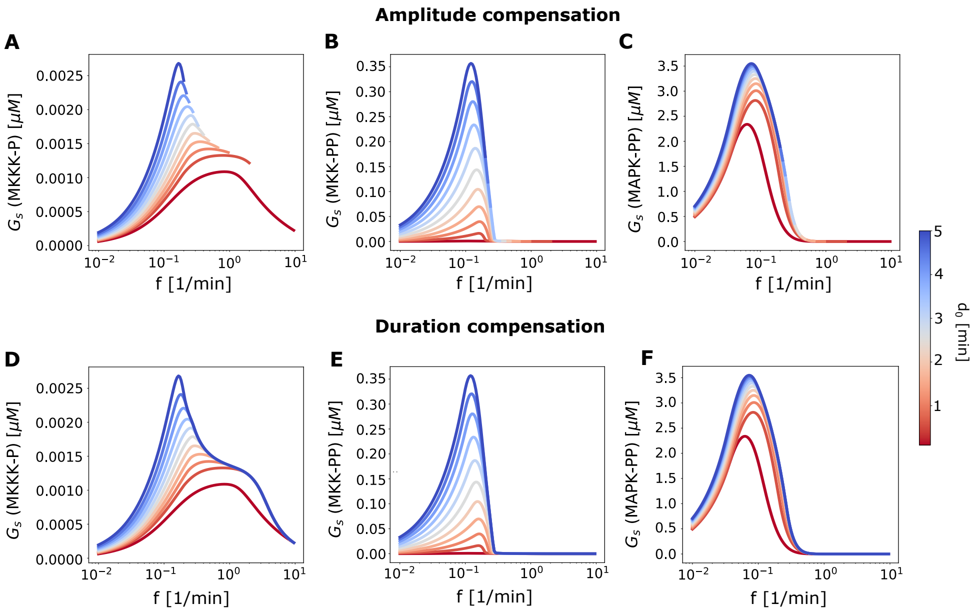


FIGURE 8. **Input's duration influence on the transient frequency response for the MAPK cascade.** Input parameters: $A_0 = 1\mu M, T_0 = 10min$. The observable $G_S$ is measured from an intial time $S = 0.1min$ in the three levels MKKK-P, MKK-PP, MAPK-PP, for the two conservation protocols (first row by amplitude compensation, second row by varying duration). The color of the curves indicates the duration of the reference input.

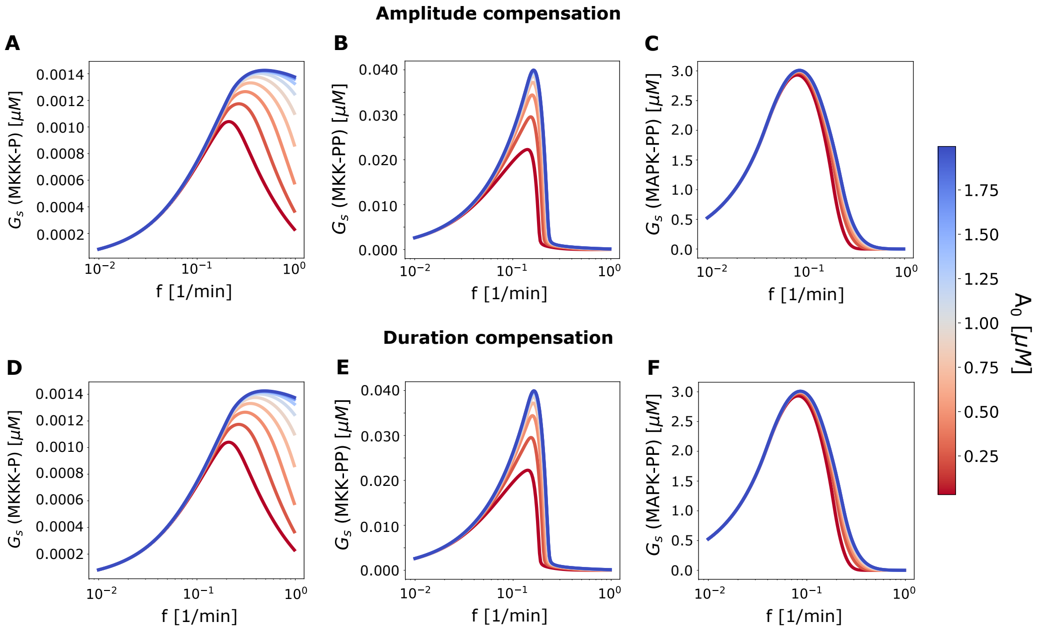


FIGURE 9. **Input's amplitude influence on the frequency response for the MAPK cascade.** Input parameters: $d_0 = 1min$, $T_0 = 10min$. The observable $G_S$ is measured from an intial time $S = 0.1min$ in the three levels MKKK-P, MKK-PP, MAPK-PP, for the two conservation protocols (first row by amplitude compensation, second row by varying duration). The color of the curves indicates the amplitude of the reference input.

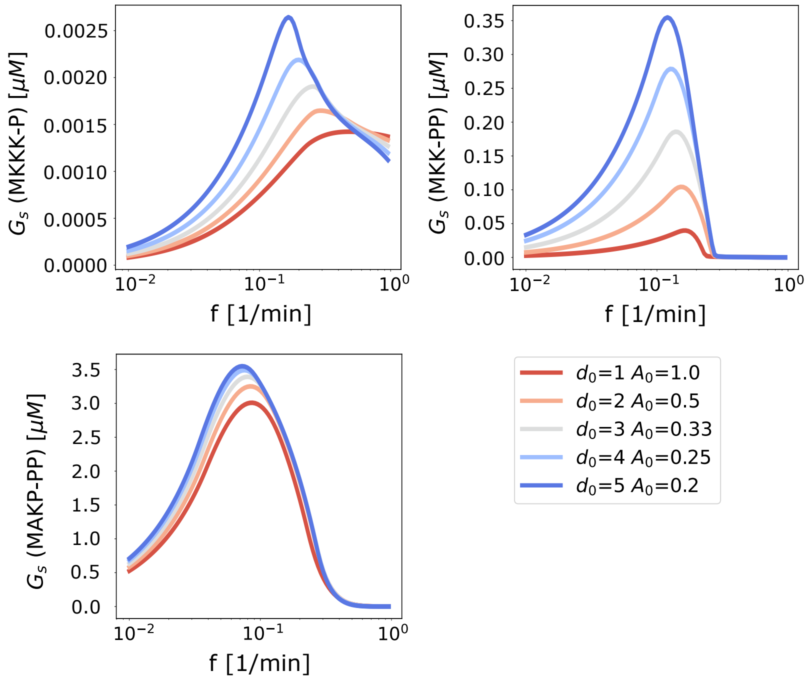


Figure 10. **Fixed input dose for different parameter inputs** ($d_0$, $A_0$).The observable $G_S$ is measured from an intial time $S = 0.1min$ in the three levels MKKK-P, MKK-PP, MAPK-PP, in the dose conservation protocol by duration compensation.
